# Repression of MAPK/Erk signaling by Efnb2-Ephb4-Rasa1 is required for lymphatic valve formation

**DOI:** 10.1101/2021.10.17.464721

**Authors:** Yaping Meng, Tong Lv, Junfeng Zhang, Anming Meng, Shunji Jia

**Affiliations:** Laboratory of Molecular Developmental Biology, State Key Laboratory of Membrane Biology, School of Life Sciences, Tsinghua University, Beijing 100084, China; Tsinghua-Peking Center for Life Sciences, Tsinghua University, Beijing 100084, China; Guangzhou Laboratory, Guangzhou 510320, Guangdong Province, China

## Abstract

The lymphatic vascular system plays important roles in various physiological and pathological processes, and lack of lymphatic or lymphovenous valves always causes lymph or blood reflux, and can lead to lymphedema. However, the molecular mechanism underlying the valve formation is poorly understood. Here we report that the MAPK/Erk signaling needs to be repressed during the valve-forming lymphatic endothelial cells (LECs) fate determination, which differs from its positive role in the LECs specification. Up-regulation of MAPK/Erk signaling in *ephb4b*, *efnb2a;efnb2b* and *rasa1a;rasa1b* mutants leads to lymphatic valve defects, whereas simultaneous loss of Erk1 and Erk2 causes valve hyperplasia. Moreover, valve defects in *ephb4b* or *rasa1a;rasa1b* mutants are mitigated in the presence of MEK inhibitors, indicating a new function of Efnb2-Ephb4-Rasa1 cassette in lymphatic valve progenitor cells specification by repressing MAPK/Erk activity. Therefore, our findings provide a mechanistic understanding of the lymphatic valve formation and potential drug targets for related lymphatic diseases.

## Introduction

The lymphatic vessel is a part of the lymphatic system and plays essential roles not only in immune cell trafficking, tissue fluid homeostasis and lipid absorption, but in many pathological conditions, including cancer progression and metastasis, lymphedema, immune responses, obesity, cardiovascular pathologies, glaucoma and neurological diseases, and so on (Koltowska, Betterman, Harvey, & Hogan, 2013; Oliver, Kipnis, Randolph, & Harvey, 2020; Schulte-Merker, Sabine, & Petrova, 2011; Tammela & Alitalo, 2010; Venero Galanternik, Stratman, Jung, Butler, & Weinstein, 2016). Lymphatic development starts with budding of lymphatic endothelial cell (LEC) progenitors expressing Prox1 from the cardinal veins or other LEC sources (Oliver, 2004; Oliver et al., 2020; Sabin, 1902; Wigle & Oliver, 1999). Once fluid flow within the lymphatic network is initiated, the expression of key transcription factors for valve development such as Gata2, Foxc2 and Prox1 is upregulated in valve-forming LECs (Geng, Cha, Mahamud, & Srinivasan, 2017; Janardhan & Trivedi, 2019; Koltowska et al., 2013). Lymphatic vessels contain one-way valves, including intraluminal lymphatic valves (LVs) and lymphovenous valves (LVVs), to ensure the unidirectional flow of lymphatic fluid. Morphological defects in these valves always compromise the maintenance of normal fluid homeostasis and result in lymphedema (Geng et al., 2017; 2016; Scallan et al., 2021). By identifying a bicuspid valve structure similar to that found in mammals, a recent study provided compelling evidence for the existence of LVs and LVVs in zebrafish facial lymphatic vessels (FLVs) (Shin et al., 2019).

Mitogen activated protein kinases (MAPKs) including ERK, JNK and p38, play important roles in a variety of biological processes, such as cell growth, migration, proliferation, differentiation, apoptosis and so on (Krens, Spaink, & Snaar-Jagalska, 2006; Kyriakis & Avruch, 2012; Mebratu & Tesfaigzi, 2009; ZHANG & LIU, 2002). Among them, MAPK/Erk pathway mainly begins with the activation of Raf by GTP-bound Ras, and then Raf phosphorylates Mek, which phosphorylates Erks, the key downstream components of the Ras-Raf-Mek-Erk signaling cascade (Fang & Richardson, 2005; Seger & Krebs, 1995). Vegfc-Vegfr3-mediated activation of MAPK/Erk signaling is the major signaling axis maintaining *Prox1* expression in LEC progenitors, and therefore promoting LEC proliferation and cell fate specification (Bui & Hong, 2020; Srinivasan et al., 2014; P. Yu, Tung, & Simons, 2014). Many studies demonstrate that Erk pathway is vital to lymphatic development. Zebrafish embryos bearing a deletion of Vegfr3 failed to initiate sprouting or differentiation of lymphatic vessels (Shin et al., 2016). Transgenic expression of RAF1^S259A^, a gain-of-function RAF1 mutant associating with human Noonan syndrome, in endothelial cells of mouse embryos activates Erk, leading to increased commitment of venous ECs to the lymphatic fate and subsequent lymphangiectasia (Deng, Atri, Eichmann, & Simons, 2013). Overexpression of Ras in the mouse endothelial cell lineage also leads to lymphatic vessel hyperplasia (T. Ichise, Yoshida, & Ichise, 2010). In addition, treatment with MEK inhibitors alleviates lymphatic anomaly-associated clinic symptom in a patient carrying an ARAF^S214P^ gain-of-function mutation, and prevents increased LEC sprouting in human primary dermal lymphatic endothelial cells (HDLECs) or zebrafish transgenic larvae that overexpress ARAF^S214P^ (Li et al., 2019). Although function of MAPK/Erk signaling in lymphatic vessel formation is well known, little is known about its role in lymphatic valve formation. One report suggests that termination of Vegfr3 signaling in collecting lymphatic trunks by Epsin1/2 is required for normal lymphatic valve development in mice, but underlying mechanism is not investigated (Xiaolei Liu et al., 2014).

Ephrin-Eph signaling mainly functions in attractive and repulsive processes, particularly in tissue boundary formation, axonal guidance, angiogenesis, and lymphangiogenesis, through guiding cell adhesion, migration, and repulsion (Arvanitis & Davy, 2008; Defourny, 2019; Klein, 2012; Liang, Patel, Janes, Murphy, & Lucet, 2019). Membrane-bound ligand Efnb2 and its tyrosine kinase receptor Ephb4 have been reported to be essential for vascular arterial-venous specification, angiogenic remodeling, and embryonic survival in mice (Barquilla & Pasquale, 2015; Hashimoto et al., 2016; Rudno-Rudzińska et al., 2017). More recently, Efnb2-Ephb4 forward signaling has been implied in regulating LV maturation during late embryonic and early postnatal development in mice (Katsuta et al., 2013; Martin-Almedina et al., 2016; Mäkinen et al., 2005; G. Zhang et al., 2015). Ras p21 protein activator 1 (Rasa1), a negative regulator of Ras through its GTPase activating protein (GAP) activity, was first discovered as a regulator of mouse embryonic growth, blood vessel formation and neuronal tissue development (Henkemeyer et al., 1995). Rasa1 can interact with Ephb4 via its non-GAP domain to facilitate its binding to and inactivation of Ras, therefore inhibiting Ras-Raf-Mek-Erk or Ras-PI3K-AKT-mTORC1 signaling in endothelial cells to regulate vascular development (Duran et al., 2019; Maertens & Cichowski, 2014). The mouse model with *Rasa1* deficiency in LECs exhibits lymphatic vessel overgrowth at the adult stage and LV endothelial cell death at the embryonic stage (Lapinski et al., 2012; 2017). Moreover, human mutations in the EFNB2-EPHB4-RASA1 cassette are related to both vascular and lymphatic diseases, such as capillary malformation-arteriovenous malformation (CM-AVM), vein of Galen malformation (VOGM), and central conducting lymphatic anomaly (CCLA) (Burrows et al., 2013; Duran et al., 2019; Eerola et al., 2003; Li et al., 2018; Martin-Almedina et al., 2016; Zeng et al., 2019). The embryonic lethality with severe cardiovascular defects in mouse *Efnb2* or *Ephb4* mutants makes it difficult to investigate Efnb2- Ephb4 signaling function in valve-forming LECs specification and to identify its downstream effector.

We previously show that zebrafish *ephb4b^tsu25^* mutants display disruption of left-right asymmetry during early embryogenesis (J. Zhang, Jiang, Liu, & Meng, 2016). Our subsequent observation surprisingly noticed a severe hemorrhage-like phenotype in *ephb4b^tsu25^* mutants at juvenile and adult stages, instigating this new study. This study demonstrates that Efnb2-Ephb4 signaling is critical for lymphatic valve progenitor cells specification by repressing MAPK/Erk activity through Rasa1.

## Results

### Blood filling of lymphatic vessels in *ephb4b* mutants

Genomic duplication in teleost fish (Amores et al., 1998) resulted in two *ephb4* genes in the zebrafish, *ephb4a* and *ephb4b*. In our previous study, we demonstrated that *ephb4b* is essential for embryonic left-right asymmetric development via regulation of dorsal forerunner cell cluster formation (J. Zhang et al., 2016). Unexpectedly, *ephb4b^tsu25^* mutants with a 25-bp deletion in the 3^rd^ exon (Figure 1A) exhibited a severe hemorrhage-like phenotype at juvenile and young adult stages, sometimes accompanied by scale protrusion and hydrops in the abdomen (Figure 1B and C). With growing old, these defects became severer and occurred in almost all of the mutants after 23 months postfertilization (mpf) (Figure 1D). By generating and observing *ephb4b^tsu25^* mutants in *Tg(lyve1b:TopazYFP)* or *Tg(lyve1b:TopazYFP;gata1:DsRed)* transgenic background, we found that the hemorrhage-like phenotype of mutants was caused by the existence of *gata1:DsRed*-positive red blood cells (RBCs) in some *lyve1b:TopazYFP*-positive lymphatic vessels (Figure 1E and F). Differing from fast circulating RBCs inside blood vessels, RBCs inside lymphatic vessels moved very slowly (Video 1).

**Figure 1.**
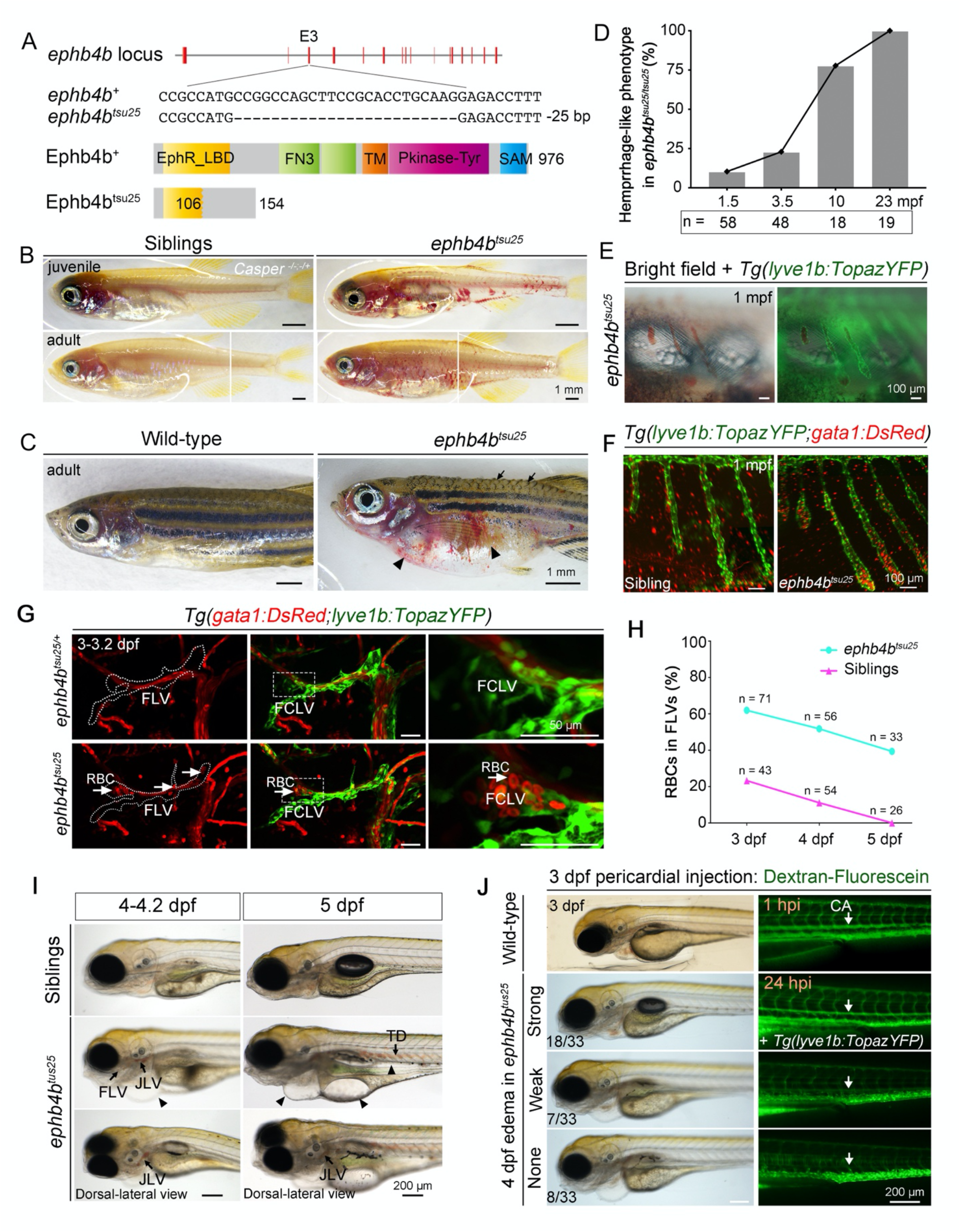
*ephb4b^tsu25^* mutant shows blood-filling of lymphatic vessels. (**A**) Generation of the *ephb4b* mutant using CRISPR/Cas9 technology with a target site at exon 3. The *ephb4b^tsu25^* mutant bears a 25-bp deletion with a protein length of 154 aa. The amino acid sequence changes after 106 aa. Refer to (J. Zhang et al., 2016) for further information. (**B**) *ephb4b^tsu25^* mutants exhibit a hemorrhage-like phenotype at the juvenile and adult stages. *ephb4b^tsu25^* mutants were crossed with *Casper* mutants (*roy^-/-^;nacre^-/-^*), and only homo-mutants for *nacre^-/-^* were selected. Scale bars, 1 mm. (**C**) A typical *ephb4b^tsu25^* adult fish in the Tubingen background shows a hemorrhage-like phenotype with edema in the heart and abdomen (arrowheads), and protrusion of scales (arrows). Scale bars, 1 mm. (**D**) Quantification of the hemorrhage-like phenotype in *ephb4b^tsu25^* mutants with embryonic development in (B-C). n = the number of fish analyzed. (**E**) In juvenile *ephb4b^tsu25^* mutants, the blood accumulates in *lyve1b:TopazYFP* labeled lymphatic vessels (green). Scale bars, 100 μm. (**F**) *gata1:DsRed*- expressing red blood cells (RBCs) accumulate in lymphatic vessels with *lyve1b:TopazYFP* expression in *ephb4b^tsu25^* larvae at 1 mpf. Scale bars, 100 μm. (**G**) *gata1:DsRed* labeled RBCs (arrows) enter the *lyve1b:TopazYFP* labeled facial lymphatic vessels (green) at 3-3.2 dpf in *ephb4b^tsu25^* mutants. Dashed lines indicate the facial lymphatic vessels (FLVs). The third panels are enlarged regions representing the facial collecting lymphatic vessels (FCLVs, boxed region). Scale bars, 50 μm. (**H**) Statistical data for the phenotype of the RBCs entering FLVs in (G). Siblings and *ephb4b^tsu25^* mutant larvae were observed at 3 dpf, 4 dpf, and 5 dpf. n = the number of fish analyzed by confocal microscopy. (**I**) Pericardial and gut edema with blood-filling of lymphatic vessels phenotype in *ephb4b^tsu25^* mutants. Arrowheads indicated the transparent edema. Arrows indicated the blood-filling in FLVs, jugular lymphatic vessels (JLVs), and the thoracic duct (TD). Scale bars, 200 μm. (**J**) Defective liquid absorption of the FLVs in *ephb4b^tsu25^* mutants. Wild-type and *ephb4b^tsu25^* mutant larvae in *Tg(lyve1b:TopazYFP)* background were injected with 2,000 kDa Dextran-Fluorescein into the pericardical region at 3 dpf, and the dispersion of the fluorescent dye into the circulation system were detected after 1 hpi and 24 hpi respectively. The ratio of embryos with exhibited pattern of the fluorescent dye is indicated. The vein is labeled by *lyve1b:TopazYFP*, while the artery is *lyve1b:TopazYFP* negative. Arrows indicate the cardinal artery (CA). Scale bars, 200 μm.

Confocal imaging of *Tg(gata1:DsRed;lyve1b:TopazYFP)* double transgenic fish showed that, in 62% (n = 44/71) of *ephb4b^tsu25^* mutant larvae, *gata1:DsRed*-positive RBCs entered *lyve1b:TopazYFP*-positive FLVs as early as 3 days postfertilization (dpf) (Figure 1G and 1H) when FLVs were just formed. In comparison, only 23% (n = 10/43) of wild-type/heterozygous sibling larvae had RBCs in FLVs at the same stage. Remarkably, 39% (n = 13/33) of mutant larvae still carried RBCs in FLVs at 5 dpf, even in the jugular lymphatic vessels (JLVs) and the thoracic duct (TD), whereas none of sibling larvae had RBCs in these lymphatic vessels (n = 0/26) at 5 dpf (Figure 1G-I; Figure 1–figure supplement 1). Accompanying blood filling in lymphatic vessels, pericardial edema was obvious in *ephb4b^tsu25^* mutants starting from 4 dpf, which sometimes expanded to the gut region (Figure 1I). Injection of 2,000 kDa Dextran-Fluorescein into the pericardial cavity of wild-type larvae in *Tg(lyve1b:TopazYFP)* background at 3 dpf led to a rapid appearance of the fluorescent dye within the artery blood vessels only after 1 hour post injection (hpi), revealing a rapid dye absorption of the lymphatic system and then subsequent transport to the circulation system. However, when this Dextran-Fluorescein was injected into the *ephb4b^tsu25^* mutant larvae in the same way, only 54.5% (n = 18/33) of embryos showed a similar dye flow pattern even at 24 hpi, whereas the other embryos only exhibited a very mild or even no absorption of the dye (Figure 1J). More importantly, this defective lymphatic absorption in *ephb4b^tsu25^* mutants was positively corelated to the edema phenotype, which was probably caused by dysfunction of the blood-filled lymphatic vessels.

In addition, we also established the *ephb4a^tsu37^* mutant line by targeting the ligand binding domain (LBD) of *ephb4a* using CRISPR/Cas9 technology, which resulted in a putative truncated protein of only 61 amino acids (Figure 1–figure supplement 2A). Unlike the *ephb4a^sa11431^* mutant embryos with a point nonsense mutation (p.Y67X) that had no phenotype in the caudal plexus (Li et al., 2018), 31.5% (n = 158/501) of *ephb4a^tsu37^* mutant embryos in *Tg(flk:mCherry;lyve1b:TopazYFP)* transgenic background showed obvious defects in caudal plexus formation at 2 dpf, which was probably caused by the failure of venous endothelial cells (ECs) separation from arterial Ecs (Figure 1–figure supplement 2B). Nevertheless, none of the *ephb4a^tsu37^* mutants had RBCs in the lymphatic vessels as did the *ephb4b^tsu25^* mutants. These results further indicate that *ephb4a* and *ephb4b* may function separately in regulating blood vessel formation and lymphatic system development, thus providing us an opportunity to investigate the roles of Ephb4 in lymphatic development in *ephb4b^tsu25^* mutants without affecting blood vessel functions.

### *ephb4b* is required for LV and LVV formation

In *ephb4b^tsu25^* mutant larvae with the *Tg(gata2:EGFP)^la3^* or *Tg(lyve1b:TopazYFP)* background, we found that the structure of *gata2a:EGFP* labeled facial LVs and FCLV-PHS LVVs that connect FCLV with primary head sinus (PHS) were defective, and sometimes even absent (Figure 2A), whereas the formation of *lyve1b:TopazYFP* labeled lymphatic vessels or blood vessels was not affected obviously compared to siblings (Figure 1G; Figure 2–figure supplement 1A). Based on the valve morphology at 3.2-3.5 dpf in *ephb4b^tsu25^* mutants, we classified the facial LVs into three types: L1, normal LV with two leaflets; L2, small LV; and L3, little or no LV structure. It was found that, when 29.7% (n = 11/37) of sibling embryos were starting to form LV structure (L1 and L2) at 3.2-3.5 dpf, 7.3% (n = 3/41) of *ephb4b^tsu25^* mutants only formed small LVs (L2) (Figure 2B and C). Similarly, the FCLV-PHS LVVs in *ephb4b^tsu25^* mutants at 3.2-3.5 dpf were categorized into three types based on their morphology: H1, normal LVV; H2, defective LVV with hollow; and H3, little or no LVV structure (Figure 2B). Our statistical data showed that, when 91.9% (n = 34/37) of sibling embryos had formed intact LVV structure (H1) at 3.2-3.5 dpf, only 41.5% (n = 17/41) of *ephb4b^tsu25^* mutant embryos had well-formed LVVs while the other mutants retained various deformed FCLV-PHS LVVs (H2 and H3) (Figure 2B and C). The FCLV-PHS LVV defects in *ephb4b^tsu25^* mutants could also be observed in the *Tg(gata2:EGFP;lyve1b:DsRed2)* background at later stages (10 dpf and 26 dpf) (Figure 2–figure supplement 1B).

**Figure 2.**
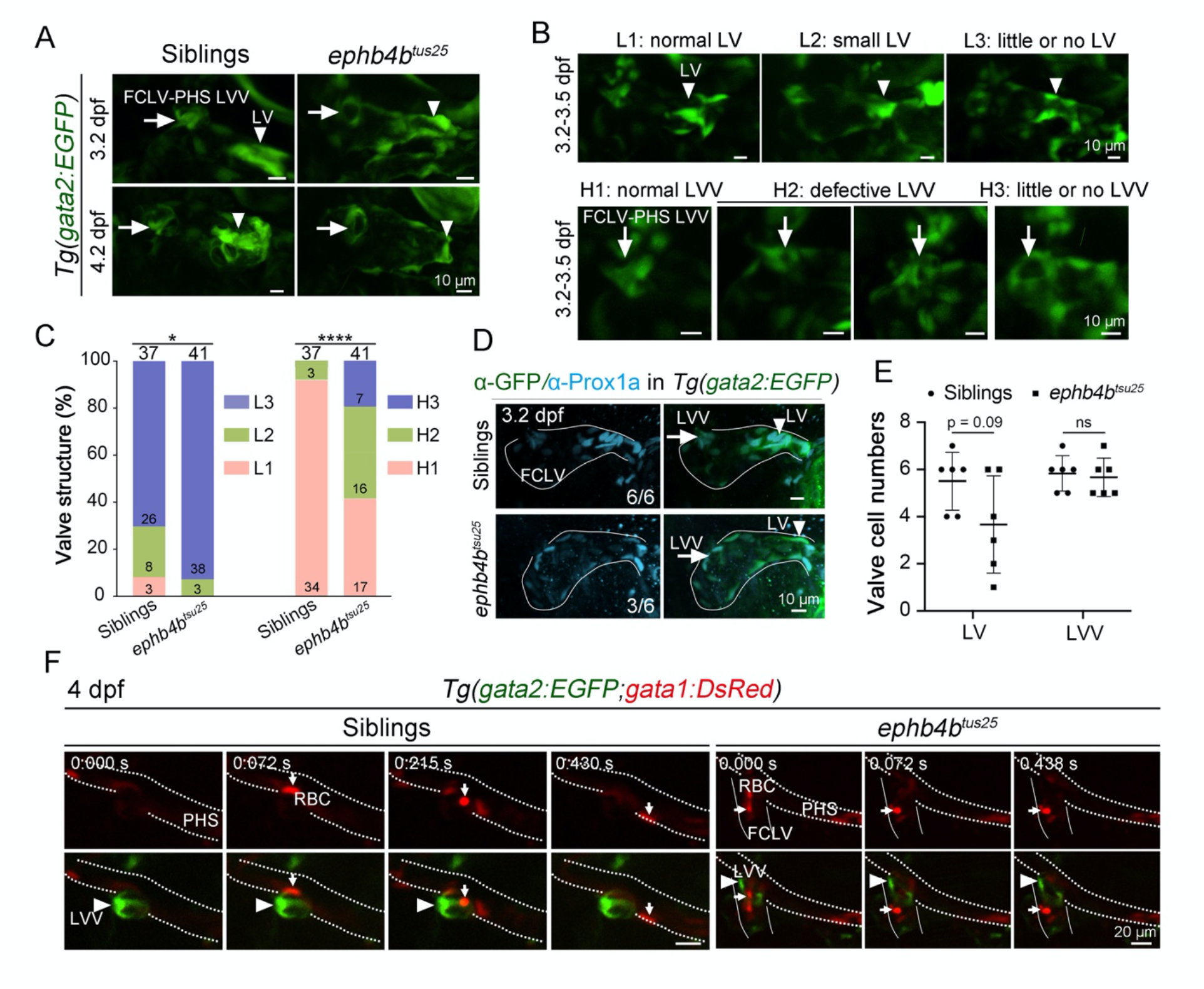
*ephb4b* is required for lymphatic valve (LV) and lymphovenous valve (LVV) formation. (**A**) *ephb4b^tsu25^* mutants show defective formation of the LVs (arrowheads) and FCLV-PHS LVVs (arrows) at 3.2 dpf and 4.2 dpf. *gata2:EGFP* was used for labeling valves. Scale bars, 10 μm. (**B**) Morphological defect classification of LV and FCLV-PHS LVV structure in *ephb4b^tsu25^* mutants. Both LVs (arrowheads) and FCLV-PHS LVVs (arrows) in mutants at 3.2-3.5 dpf can be divided into 3 categories depending on their morphological integrity, including L1-L3 and H1-H3 respectively. Scale bars, 10 μm. (**C**) Statistical summary of the valve defects in siblings or *ephb4b^tsu25^* mutants at 3.2-3.5 dpf in (B). Chi-squared test, *, p < 0.05; ****, p < 0.0001. (**D**) Immunofluorescence results of Prox1a in siblings and *ephb4b^tsu25^* mutants in the *Tg(gata2:EGFP)* transgenic background at 3.2 dpf. The FCLV-PHS LVVs and LVs are indicated by arrows and arrowheads, respectively. The ratio of embryos with exhibited pattern of the Prox1a expression (cyan) in LV is indicated. Scale bars, 10 μm. (**E**) Statistical analysis of the *gata2:EGFP*-positive valve-forming LECs in siblings (n = 6) and *ephb4b^tsu25^* mutants (n = 6) in (D). Unpaired *t* test, ns, no statistical significance. (**F**) Confocal imaging of RBCs movement near the FCLV-PHS LVV in siblings and *ephb4b^tsu25^* mutants in the *Tg(gata2:EGFP;gata1:DsRed)* background. Primary head sinus (PHS) and FCLV are marked with white dotted lines and solid lines respectively. RBCs (arrows) are labeled by *gata1:DsRed*, and FCLV-PHS LVVs (arrowheads) are *gata2:EGFP* positive. RBCs enters the FCLV through the defective LVV in *ephb4b^tsu25^* mutants, but are blocked by the well-formed LVV in siblings at 3.2 dpf. Scale bars, 20 μm.

It is reported that high-level *Prox1* expression happens in valve-forming LECs, which further cluster together and change the nuclear direction perpendicular to lymph flow to construct valve structure (Geng et al., 2017; Janardhan & Trivedi, 2019). Our immunofluorescence results in siblings in the *Tg(gata2:EGFP)* transgenic background showed that the expression of Prox1a, the zebrafish homolog of the mammalian Prox1, together with *gata2:EGFP,* was much higher in valve-forming LECs in the LV and LVV structure at 3.2 dpf. In *ephb4b^tsu25^* mutants, however, the number of valve-forming LECs with high-level Prox1a in the LVs was reduced to some extent (Figure 2D and E). These results indicate that *ephb4b* might be involved in the initiation of the lymphatic valve formation.

Then, confocal live imaging was applied to record the blood flow near the FCLV-PHS LVV in the *Tg(gata2:EGFP;gata1:DsRed)* background. In siblings, *gata1:DsRed-*expressing RBCs in PHS were blocked from flowing into the FCLV by *gata2:EGFP-*expressing LVV at 4 dpf (n = 7; Figure 2F; Video 2). In *ephb4b^tsu25^* mutants, however, RBCs in the PHS were able to flow into the FCLV via the intervening space of the defective FCLV-PHS LVV (n = 6; Figure 2F; Video 2). This observation supports the idea that *ephb4b* is essential for valve formation in the lymphatic system.

### *efnb2* is required for LV and LVV formation

Efnb2-Ephb4 forward signaling has been demonstrated to regulate the LV maturation in mice (G. Zhang et al., 2015). To assess the requirement of *efnb2a* and *efnb2b,* two homologues of the mammalian *Efnb2*, in lymphatic valve initiation process, we knocked out *efnb2a* and *efnb2b* individually in zebrafish using CRISPR/Cas9 technology (Figure 3A and B). Homozygous *efnb2a^tsu41^* mutants with a 7-bp deletion, which putatively express a truncated Efnb2a protein only with its receptor binding domain (RBD) with loss of the transmembrane (TM) domain (Figure 3A), exhibited caudal plexus malformation (Figure 3C), which was also seen in *ephb4a^tsu37^* mutants (Figure 2–figure supplement 1B), and blood reflux into lymphatic vessels with edema phenotype (Figure 3, D and E), which resembled the *ephb4b^tsu25^* phenotype (Figure 1I). In contrast, *efnb2b^tsu42^* mutants, which carried a 15-bp deletion and an 8-bp insertion that presumably expressed a mutant Efnb2b protein without the intact RBD (Figure 3B), developed normally (Figure 3D and E). By crossing double heterozygous fish, we obtained *efnb2a^tsu41^*;*efnb2b^tsu42^* double mutants. We observed the lymphatic blood filling and pericardial edema defects in all of the double mutants (Figure 3D), which were much severer than *efnb2a^tsu41^* single mutants, suggesting a possible compensatory effect of *efnb2b* in *efnb2a^tsu41^* mutants. Confocal imaging confirmed the absence of LVs and FCLV-PHS LVVs labeled with *gata2:EGFP* in *efnb2a^tsu41^*;*efnb2b^tsu42^* double mutants with the *Tg(gata2:EGFP;lyve1:DsRed2)* transgenic background (Figure 3F). Immunostaining results showed that, distinguished from ductal LECs that expressed medium levels of Prox1a, valve-forming LECs with high levels of Prox1a clustered at putative valve sites in sibling embryos, while these kinds of clustered valve-forming cells with high-level Prox1a were diminished in *efnb2a^tsu41^*;*efnb2b^tsu42^* double mutants, indicating a failure of these cells in valve-forming LECs fate commitment (Figure 3G and H; Video 3). Taken together, these results suggest that the Efnb2-Ephb4 pathway participates in valve-forming LEC formation, most likely in the fate specification of the progenitor cells.

**Figure 3.**
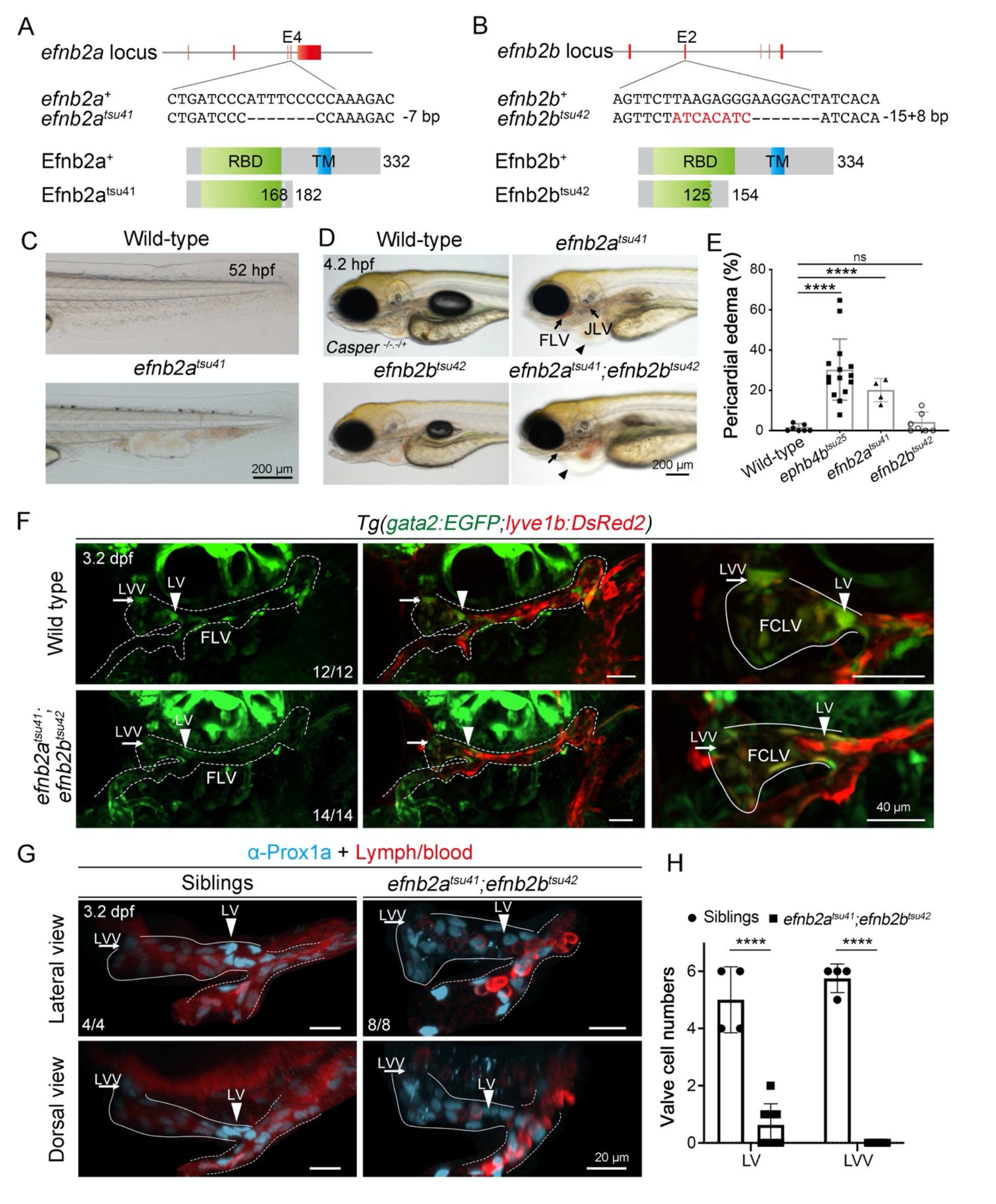
*efnb2* is required for lymphatic valve (LV) and lymphovenous valve (LVV) formation. (**A**) Generation of the *efnb2a* mutant using CRISPR/Cas9 technology with a target site at exon 4. The *efnb2a^tsu41^* mutant bears a 7-bp deletion with a protein length of 182 aa. The amino acid sequence changes after 168 aa. The *efnb2a^tsu41^* mutant protein has an intact receptor binding domain (RBD). (**B**) Generation of the *efnb2b* mutant using CRISPR/Cas9 technology with a target site at exon 2. The *efnb2b^tsu42^* mutant bears a 15-bp deletion and an 8-bp insertion with a protein length of 154 aa. The amino acid sequence changes after 125 aa. (**C**) *efnb2a^tsu41^* mutants exhibit abnormal blood vessel formation and blood accumulation in the tail region at 52 hpf, similar to *ephb4a^tsu37^* mutants (refer to Figure S2B). Scale bar, 200 μm. (**D**) *efnb2a^tsu41^* and *efnb2a^tsu41^;efnb2b^tsu42^* double mutants exhibit pericardial edema (arrowheads) and blood-filling of facial lymphatic vessels (arrows) at 4.2 dpf, while *efnb2b^tsu42^* develops normally. Scale bar, 200 μm. (**E**) Statistical analysis of the pericardial edema phenotype in *ephb4b^tsu25^*, *efnb2a^tsu41^* and *efnb2b^tsu42^* mutants. Unpaired *t* test, ****, p < 0.0001. Each dot represents the percentage of edema embryos (with total embryos > 40) from one pair of fish with the indicated genotype. (**F**) No valve structure formation in *efnb2a^tsu41^;efnb2b^tsu42^* double mutants at 3.2 dpf. The facial lymphatic vessels (FLVs) and facial collecting lymphatic vessel (FCLV) are marked by dotted lines and solid lines respectively. LVs and FCLV-PHS LVVs are indicated by arrowheads and arrows, respectively. The ratio of embryos with exhibited valve structure is indicated. Lateral views, anterior to the left. Scale bars, 40 μm. (**G**) Immunofluorescence results of Prox1a in siblings and *efnb2a^tsu41^;efnb2b^tsu42^* mutants at 3.2 dpf. FCLV-PHS LVVs and LVs are indicated by arrows and arrowheads, respectively. The ratio of embryos with exhibited pattern of the Prox1a expression (cyan) is indicated. The blood or lymph autofluorescence can be detected as red fluorescence. Siblings were defined as neither *efnb2a^tsu41^* mutant nor *efnb2b^tsu42^* mutant. Lateral or dorsal views are shown, and anterior to the left. Scale bars, 20 μm. (**H**) Statistical analysis of the high-level Prox1a-positive valve-forming LECs in siblings (n = 4) and *efnb2a^tsu41^;efnb2b^tsu42^* mutants (n = 8) in (G). Unpaired *t* test, ****, p < 0.0001.

### *Rasa1* is required for LV and LVV formation

Efnb2-Ephb4 forward signaling can activate various downstream effectors, including Src, Rac, Fak, RhoA, Crk, Nck, Abl, Rasa1, and Grb2, among which *Rasa1* mutation has been found to be involved in LV dysfunction (Lapinski et al., 2017; Shiuan & Chen, 2016; Yang, Wei, Chen, & Wu, 2018). To investigate the function of *rasa1* in the zebrafish, we generated the *rasa1a^tsu38^* mutant line that carries a 2-bp deletion and an 18-bp insertion in the first SH2 domain and the *rasa1b^tsu39^* mutant line with a 308-bp deletion and a 31-bp insertion near the start codon (Figure 4A and B). *rasa1a^tsu38^* or *rasa1b^tsu39^* homozygous mutants developed normally to adulthood without visible morphological changes. However, *rasa1a^tsu38^;rasa1b^tsu39^* double mutants exhibited blood filling within lymphatic vessels and severe pericardial edema at 4 dpf, and could not survive over 6 dpf (Figure 4C). In addition, we obtained another *rasa1a^tsu40^* mutant allele with only a 3-bp deletion that resulted in a loss of the corresponding isoleucine (I152) in the central β-sheet, just before the arginine (R153) (Figure 4A), which is critical for SH2 domain binding to a phosphotyrosine peptide (pTyr) (Jaber Chehayeb, Stiegler, & Boggon, 2019; Pamonsinlapatham et al., 2009). We found that all of the *rasa1a^tsu40^*;*rasa1b^tsu39^* or *rasa1a^tsu38/tsu40^*;*rasa1b^tsu39^* double mutants exhibited similar pericardial edema and blood-filling lymphatic vessels (Figure 4C) as did *rasa1a^tsu38^*;*rasa1b^tsu39^* mutants, suggesting that the binding ability of the Rasa1 SH2 domain to pTyr proteins is important for its functions in lymphatic valve formation. To make live imaging possible, we obtained *rasa1a^tsu38^;rasa1b^tsu39^* double mutants in the *Tg(gata2:EGFP;gata1:DsRed), Tg(lyve1b:TopazYFP)* or *Tg(gata2:EGFP;lyve1b:DsRed2)* transgenic background. Confocal imaging revealed that the *gata2:EGFP-*expressing LVs and FCLV-PHS LVVs in FLVs were almost completely lost in *rasa1a^tsu38^;rasa1b^tsu39^* double mutants at 3-4 dpf, while the *lyve1b:TopazYFP*-expressing lymphatic vessels from facial lymphatic sprout (FLS) were over-proliferated (Figure 4D). This observation suggests that *rasa1a* and *rasa1b* cooperatively promote the lymphatic valve development but repress lymphatic vessel growth. Similarly, *gata2:EGFP-*expressing valve structures were almost absent in *rasa1a^tsu40^*;*rasa1b^tsu39^* mutants in the *Tg(gata2:EGFP;lyve1b:DsRed2)* transgenic background (Figure 4–figure supplement 1A). Additionally, we recorded the valve-forming LECs behavior by confocal imaging in the *Tg(gata2:EGFP;gata1:DsRed)* background, and found that, compared to the siblings, the valve- forming LECs with high-level Prox1a could hardly be detected in *rasa1a^tsu38^;rasa1b^tsu39^* double mutants at the putative LVV and LV sites so that no valve leaflets were formed (Figure 4E and F).

**Figure 4.**
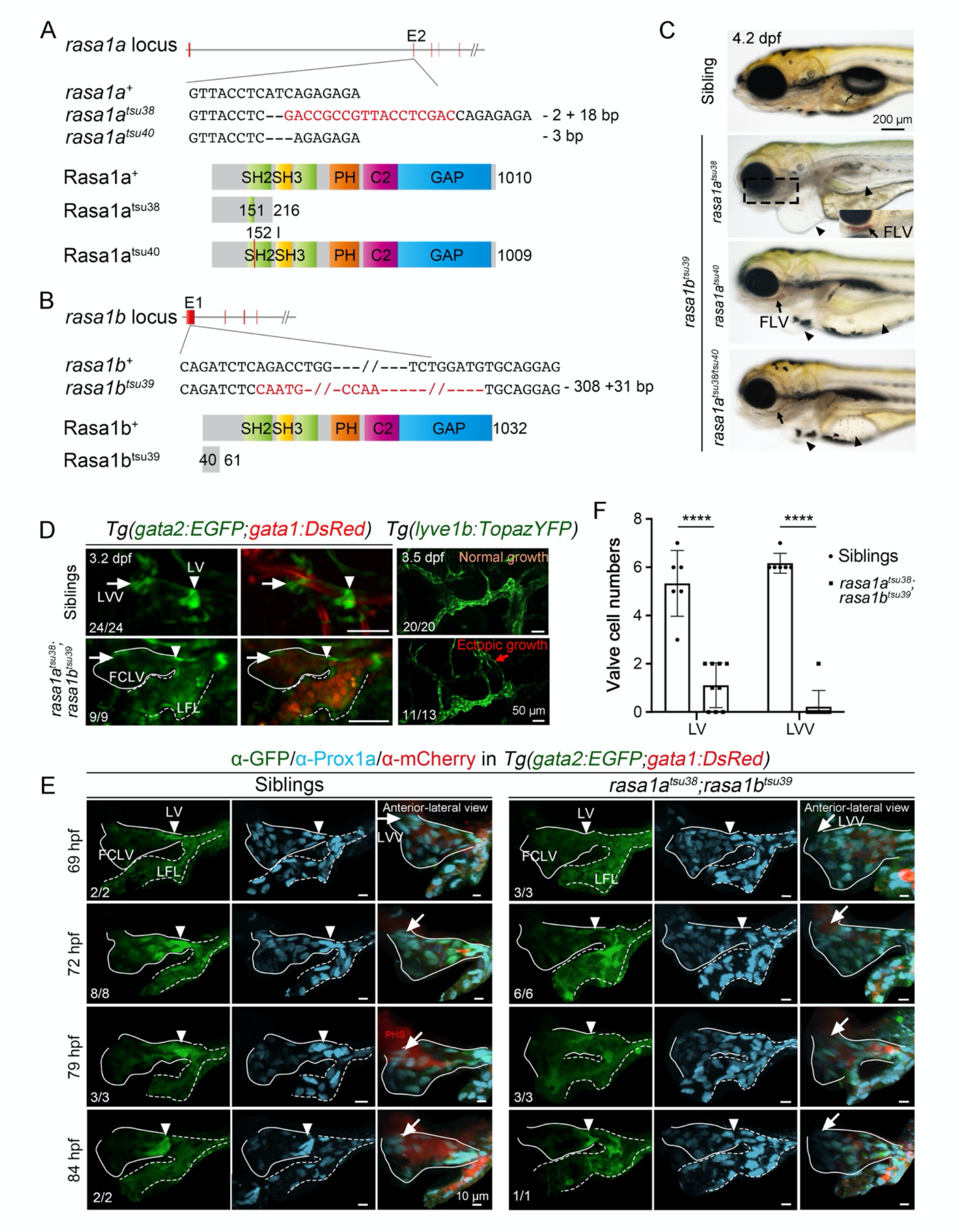
*rasa1* is required for lymphatic valve (LV) and lymphovenous valve (LVV) formation. (**A**) Generation of the *rasa1a* mutant using CRISPR/Cas9 technology with a target site at exon 2. Two mutant alleles were generated. The *rasa1a^tsu38^* mutant bears a 2-bp deletion and 18-bp insertion with a protein length of 216 aa. The amino acid sequence changes after 151 aa. The *rasa1a^tsu40^* mutant bears a 3-bp deletion with loss of the isoleucine (I) at 152 aa. (**B**) Generation of the *rasa1b* mutant using CRISPR/Cas9 technology with a target site at exon 1. The *rasa1b^tsu39^* mutant bears a 308-bp deletion and 31-bp insertion with a protein length of 61 aa. The amino acid sequence changes after 40 aa. (**C**) Double mutants of *rasa1a* and *rasa1b* exhibit pericardial edema (arrowheads) and blood-filling of facial lymphatic vessels (arrows) at 4.2 dpf. Note that the compound heterozygote of *rasa1a^tsu38/tsu40^* with the *rasa1b^tsu39^* mutant also exhibits a similar phenotype. Lateral-ventral view of the boxed region is shown at the right corner. Mutants with Tubingen background were used. Scale bar, 200 μm. (**D**) Severe valve malformation in *rasa1a^tsu38^;rasa1b^tsu39^* double mutants at 3.2 dpf. LVs and FCLV-PHS LVVs or their expected positions are indicated by arrowheads and arrows, respectively. The FCLV and lateral facial lymphatic vessel (LFL) in mutants are marked by solid lines and dotted lines respectively. Note the existence of RBCs labeled by *gata1:DsRed* in the FLVs, and the overgrowth of the *lyve1b:TopazYFP* labeled facial lymphatic sprout (FLS) in *rasa1a^tsu38^;rasa1b^tsu39^* mutants. The ratio of embryos with exhibited phenotype is indicated. Siblings were defined as neither *rasa1a^tsu38^* mutant nor *rasa1b^tsu39^* mutants. Scale bars, 50 μm. (**E**) Prox1a immunostaining (cyan) in siblings and *rasa1a^tsu38^;rasa1b^tsu39^* double mutants in the *Tg(gata2:EGFP;gata1:DsRed*) background. The high Prox1 signal (cyan) reveals the specification of valve-forming LECs. In siblings, FCLV-PHS LVVs (arrows) and LVs (arrowheads) start forming from about 69 hpf and 72 hpf, respectively. No valve specification in *rasa1a^tsu38^;rasa1b^tsu39^* mutants. The third panels are the anterior-lateral view for observing the FCLV-PHS LVVs. The ratio of embryos with exhibited pattern of the Prox1a expression (cyan) is indicated. Siblings defined as neither *rasa1a^tsu38^* mutant nor *rasa1b^tsu39^* mutants. Scale bars, 10 μm. (**F**) Statistical analysis of the *gata2:EGFP*-positive valve-forming LECs in siblings (n = 9) and *rasa1a^tsu38^;rasa1b^tsu39^* mutants (n = 6) at 3.2-3.5 dpf in (E). Unpaired *t* test, ****, p < 0.0001.

### Simultaneous loss of Erk1 and Erk2 leads to valve hyperplasia

The above data indicate that Efnb2-Ephb4-Rasa1 regulates the development of both the LV and LVV formation in the zebrafish, but how they function is still unclear. It is reported that Rasa1 can directly bind to pTyr of Ephb4 and consequently inactivates the Ras-Raf-Mek-Erk or Ras- PI3K-AKT-mTORC1 pathway by switching the active GTP-bound Ras to the inactive GDP-bound form (Haupaix et al., 2013; Xiao et al., 2012; Zeng et al., 2019). Given that treatment with the MEK inhibitors leads to an abrupt improvement in symptoms of lymphatic disorders (Li et al., 2019), we speculated that Efnb2-Ephb4 signaling might regulate valve formation by inhibiting the Ras-Raf-Mek-Erk pathway via Rasa1.

To explore the roles of MAPK/Erk signaling in lymphatic valve formation, we generated the *erk1^tsu45^* mutant line with a 47-bp deletion and the *erk2^tsu46^* mutant line with a 5-bp deletion near the start codon (Figure 5A and B). Likely due to compensatory function of Erk1 and Erk2, either *erk1^tsu45^* or *erk2^tsu46^* homozygous mutants developed normally to adulthood. We then generated *erk1^tsu45^;erk2^tsu46^* zygotic double mutants to observe potential phenotypic changes. Pericardial edema was obvious in 70% (n = 19/27) of *erk1^tsu45^;erk2^tsu46^* mutants at 5 dpf and sometimes expanded to the gut region (Figure 5C and D). To observe lymphatic valve formation, *erk1^tsu45^;erk2^tsu46^* double mutants were introduced into the *Tg(lyve1b:DsRed2;gata2:EGFP)* background and subjected to confocal microscopy. We observed that, compared to *erk1^tsu45/+^*;*erk2^tsu46/+^* double heterozygote larvae, the *gata2:EGFP*-positive valve-forming LECs were increased obviously in *erk1^tsu45^;erk2^tsu46^* double mutants (Figure 5E). On the other hand, compared to morphology of the *lyve1b:DsRed2-*labeled FLVs in *erk1^tsu45/+^*;*erk2^tsu46/+^* double heterozygote larvae at 5 dpf, the anterior part of LFL and the otolithic lymphatic vessels (OLV) in *erk1^tsu45^;erk2^tsu46^* double mutants were defective or completely lost during the FLV formation (Figure 5E and F), which is consistent with the previous report that Erk inhibition blocks trunk lymphatic sprouting and differentiation (Shin et al., 2016). These data indicate that MAPK/Erk signaling promotes lymphatic vessel formation, but represses the LV and LVV development.

**Figure 5.**
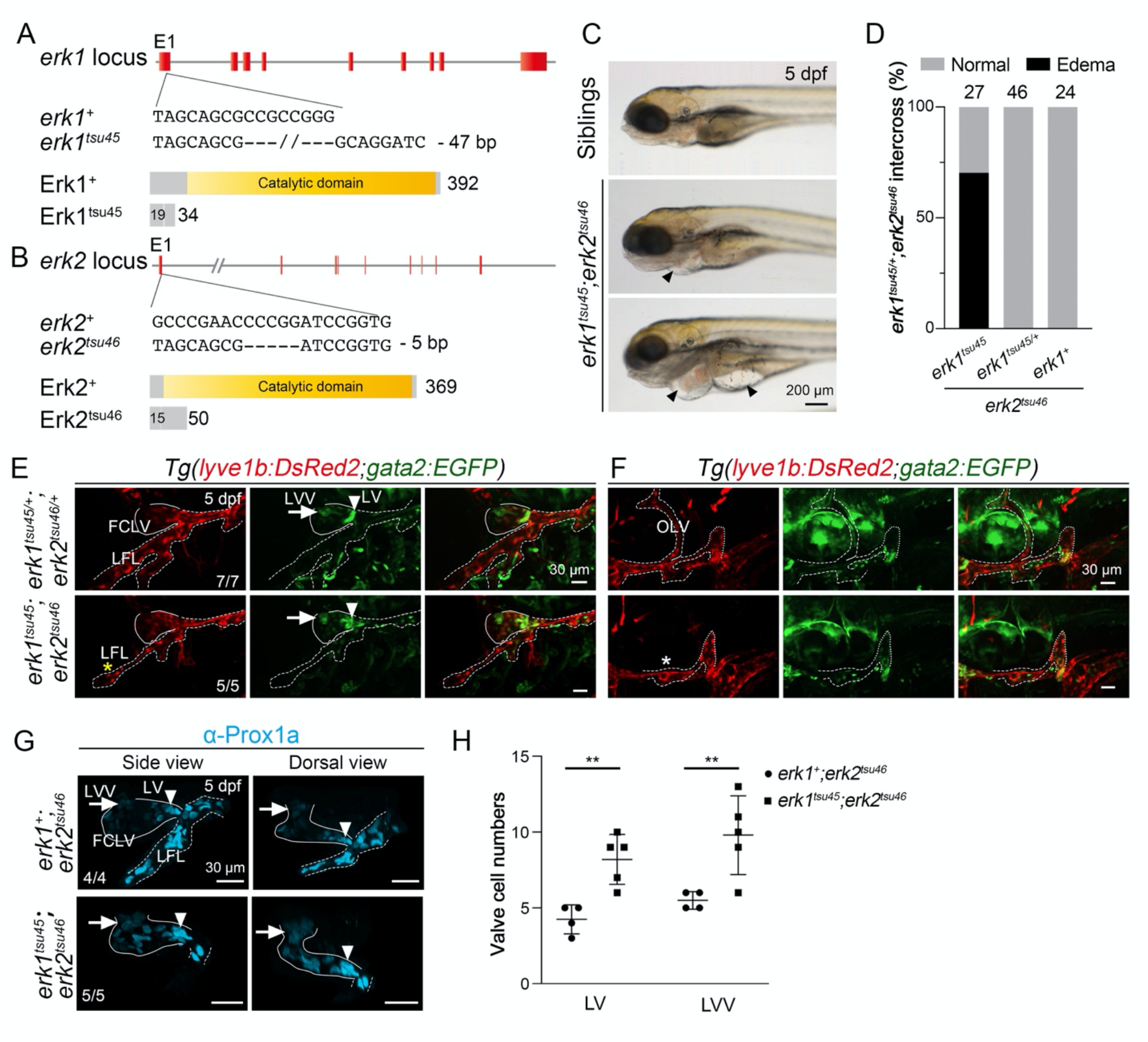
*erk1/2* inhibits valve formation in zebrafish lymphatic system. (**A**) Generation of the *erk1* mutant using CRISPR/Cas9 technology with a target site at exon 1. The *erk1^tsu45^* mutant bears a 47-bp deletion with a protein length of 34 aa. The amino acid sequence changes after 19 aa. (**B**) Generation of the *erk2* mutant using CRISPR/Cas9 technology with a target site at exon 1. The *erk2^tsu46^* mutant bears a 5-bp deletion with a protein length of 50 aa. The amino acid sequence changes after 15 aa. (**C**) *erk1^tsu45^*;*erk2^tsu46^* double mutants exhibit slight (middle panel) or severe (bottom panel) pericardial edema at 5 dpf. Mutants were generated using *erk1^tsu45/+^*;*erk2^tsu46/+^* double heterozygotes with *Casper ^-/+;-/+^* background. Scale bar, 200 μm. (**D**) Statistical summary of pericardial edema in the offsprings from the intercrossing of *erk1^tsu45/+^*;*erk2^tsu46^* mutants. 70% (n = 19/27) of *erk1^tsu45^*;*erk2^tsu46^* double mutants exhibit pericardial edema. (**E**) Valve hyperplasia and lymphatic vessel lost in *erk1^tsu45^*;*erk2^tsu46^* double mutants. The yellow asterisks show the malformed anterior LFL. Arrows and arrowheads indicate the FCLV-PHS LVVs and LVs, respectively. Solid and dotted lines indicate the FCLV and LFL, respectively. Lateral view, anterior to the left. Scale bars, 30 μm. (**F**) The otolithic lymphatic vessel (OLV) is lost in *erk1^tsu45^*;*erk2^tsu46^* double mutants. The white asterisk shows the region where the OLV sprouts from the LFL. The dotted line indicates the facial lymphatic sprout (FLS). Lateral view, anterior to the left. Scale bars, 30 μm. (**G**) Prox1a immunostaining (cyan) reveals overgrowth of valve-forming LECs in *erk1^tsu45^*;*erk2^tsu46^* double mutants. Note that the number of valve-forming LECs in LVs (arrowheads) and FCLV-PHS LVVs (arrows) are increased in *erk1^tsu45^*;*erk2^tsu46^* double mutants. Scale bars, 30 μm. (**H**) Statistical analysis of the high-level Prox1a-positive valve-forming LECs in *erk1^+^*;*erk2^tsu46^* (n=4) or *erk1^tsu45^*;*erk2^tsu46^* (n=5) mutants at 5 dpf in (G). Unpaired *t* test, **, p < 0.01.

Then, we further examined lymphatic valve formation by Prox1a immunostaining. In *erk2^tsu46^* single mutants at 5 dpf, the number of the valve-forming LEC cells with high-level Prox1a in LV and LVV were 4.25 and 5.5 on average, respectively, while these numbers increased to 8.2 and 9.8 on average in *erk1^tsu45^;erk2^tsu46^* double mutants (Figure 5G and H). These results support the idea that the inhibition of MAPK/Erk signaling results in lymphatic valve hyperplasia at early stages of lymphatic system development.

### Efnb2-Ephb4-Rasa1 signaling regulates lymphatic valve initiation by inhibiting the Ras-Raf-Mek-Erk pathway

To confirm the role of Efnb2-Ephb4-Rasa1 signaling in lymphatic valve formation by inhibiting the Ras-Raf-Mek-Erk pathway, we examined phospho-Erk1/2 (p-Erk1/2) levels by immunofluorescence in isolated *gata2:EGFP/lyve1b:DsRed2* double-positive putative valve-forming LECs at 3.2 dpf from sibling or *ephb4b^tsu25^* mutant larvae in the *Tg(gata2:EGFP;lyve1b:DsRed2)* transgenic background. Results showed that p-Erk1/2 signal was higher in *ephb4b^tsu25^* mutants than in siblings (Figure 6A and B). Considering that valve- forming LECs derive from lymphatic vessels anatomically nearby the lymphatic valve structure (Ducoli & Detmar, 2021; Geng et al., 2017; Sabine et al., 2012), we wondered whether *efnb2a/efnb2b* and *ephb4b* have specific expression patterns there. Fluorescence *in situ* hybridization (FISH) showed that both FCLVs and LFLs expressed *efnb2a* and *efnb2b*, whereas only the LFLs expressed *ephb4b* (Figure 6C; Figure 6–figure supplement 1B). When examining *in situ* p-Erk1/2 levels in sibling larvae at 3.2 dpf with immunofluorescence, we observed low levels of p-Erk1/2 in FCLV and anterior LFL (aLFL) endothelial cells, in contrast to higher levels of p-Erk1/2 in posterior LFL (pLFL) (Figure 6D). However, elevated p-Erk1/2 levels in the FCLV and aLFL endothelial cells were observed in *ephb4b^tsu25^*, *efnb2a^tsu41^;efnb2b^tsu42^* and *rasa1a^tsu38^*;*rasa1b^tsu39^* mutant larvae (Figure 6D). Taken together, these data suggest that Efnb2- Ephb4-Rasa1 signaling might act to downregulate MAPK/Erk signaling in LECs so as to allow the LV and LVV formation.

**Figure 6.**
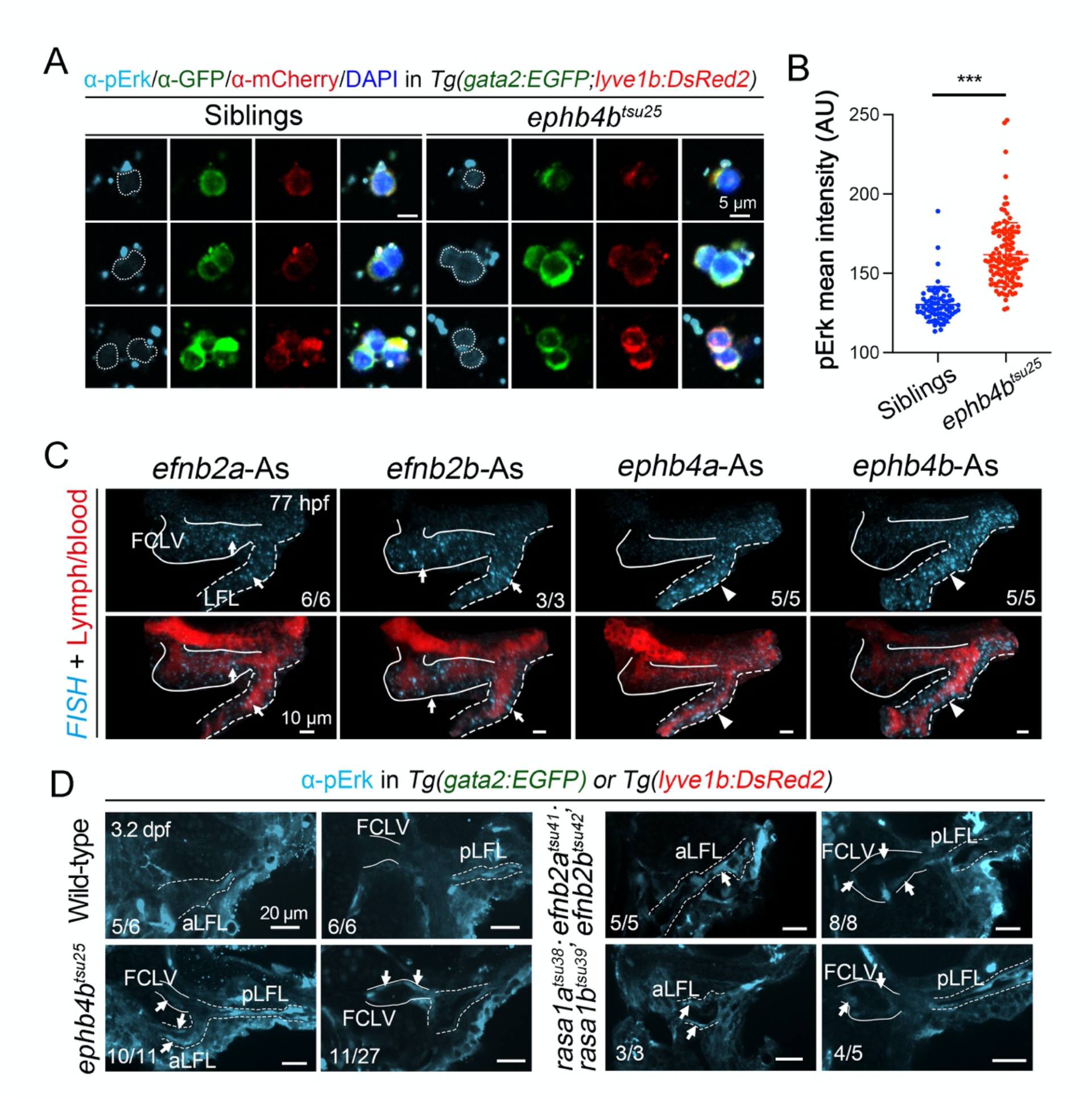
Increased pErk in some LECs in Efnb2-Ephb4-Rasa1 signaling mutants. (**A**) pErk immunostaining (cyan) of *gata2:EGFP* and *lyve1b:DsRed2* double positive cells sorted from siblings and *ephb4b^tsu25^* mutant embryos at 3.2 dpf. Scale bars, 20 μm. (**B**) Statistical results of the pErk fluorescence intensities in the single cells of siblings and *ephb4b^tsu25^* mutant in (A). All fluorescence intensities are plotted in a.u. Unpaired *t* test, ***, p < 0.001. (**C**) FISH staining of the *efnb2a*, *efnb2b*, *ephb4a,* and *ephb4b* in facial lymphatic vessels. Arrows indicate the expression of *efnb2a* and *efnb2b* in the facial collecting lymphatic vessel (FCLV, solid lines) and lateral facial lymphatic (LFL, dotted lines) at 77 hpf. Arrowheads indicate the expression of *ephb4a* and *ephb4b* in the LFL. Lateral views, anterior to the left. Scale bars, 10 μm. (**D**) Increased pErk expression in the lymphatic endothelial cells of *ephb4b^tsu25^*, *efnb2a^tsu41^;efnb2b^tsu42^*, or *rasa1a^tsu38^;rasa1b^tsu39^* mutants at 3.2 dpf. pErk expression in the FCLV and anterior LFL (aLFL) in wild-type embryos is hard to detect. The increased expression of the pErk in the FCLV (solid lines) or LFL (dotted lines) endothelial cells are marked by arrows in these mutants. The ratio of embryos with exhibited pattern of the pErk expression (cyan) is indicated. Lateral views, anterior to the left. Scale bars, 20 μm.

Next, we tried to use a set of MEK inhibitors to rescue the lymphatic valve defects in *ephb4b^tsu25^* mutants. We applied four different MEK inhibitors, including Selumetinib, Cobimetinib, Trametinib, and U0126-EtOH, at different concentrations to *ephb4b^tsu25^* mutants from 2 dpf to 4 dpf and used pericardial edema as the phenotypic readout (Figure 7A). As a result, we found that all the four MEK inhibitors could partially rescue the edema phenotype of *ephb4b^tsu25^* mutants at proper concentrations (Figure 7B). In contrast, the mTOR inhibitors rapamycin and BEZ235 could not rescue the edema phenotype of *ephb4b^tsu25^* mutants (Figure 7–figure supplement 1). Furthermore, we checked the formation of LVs and LVVs in *ephb4b^tsu25^* mutants in the *Tg(gata2:EGFP)* transgenic background after 10 μM Selumetinib treatment, and found that the *gata2:EGFP*-labeled valve structure could be restored to a certain extent at 3.2-3.5 dpf (Figure 7C). Similar rescue effects on the edema formation were also observed in *rasa1a^tsu40^;rasa1b^tsu39^* double mutants with Selumetinib treatment at 10 μM or 100 μM (Figure 7D). Moreover, immunofluorescence with anti-Prox1 antibody indicated that the Prox1a-positive valve-forming LECs could be restored in *rasa1a^tsu40^;rasa1b^tsu39^* double mutants with 100 μM Selumetinib treatment (Figure 7E and F).Therefore, we propose that Efnb2-Ephb4-Rasa1 signaling inhibits Ras-Raf-Mek-Erk activity to promote valve-forming LECs specification, therefore subsequently lymphatic valves formation.

**Figure 7.**
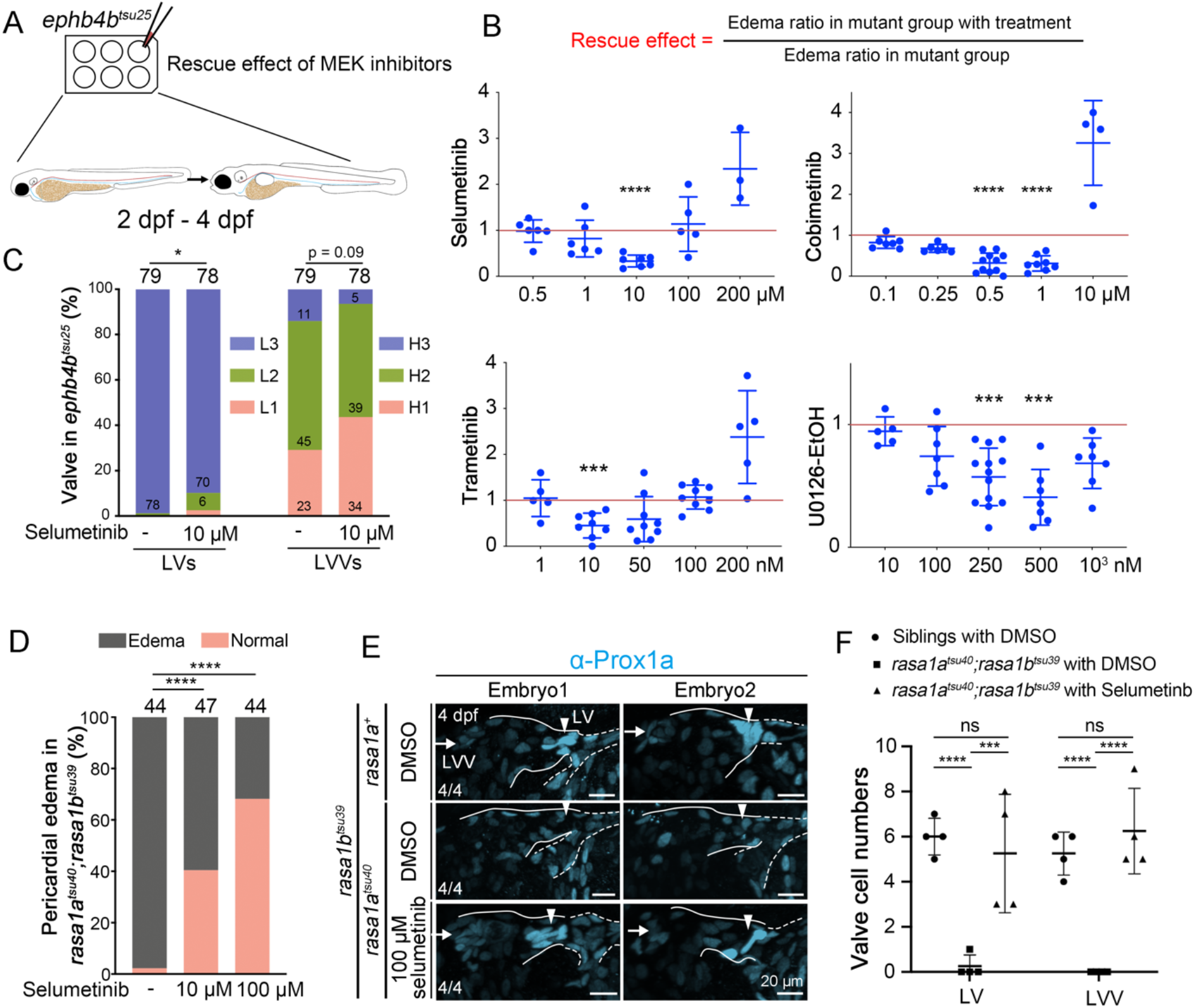
Inhibition of Mek-Erk signaling is required for valve-forming LEC specification. (**A**) Inhibitor treatment strategy of *ephb4b^tsu25^* mutants. Treatments started from 2 dpf in 6-well plates and continued until 4 dpf. The inhibitor effects were evaluated by comparing the edema ratio in mutant group with inhibitor treatment to that in the mutant group without inhibitor treatment. (**B**) MEK inhibitor treatments can decrease the edema ratio in *ephb4b^tsu25^* mutants. Selumetinib, Cobimetinib, Trametinib, and U0126-EtOH were used at proper concentrations as indicated. The vertical axis represents rescue effect value, and samples below the red lines indicate rescue effects. Chi-squared test, ***, p < 0.001; ****, p < 0.0001. (**C**) Statistical data of the Selumetinib rescue effects on LV and LVV structure in *ephb4b^tsu25^* mutants at 3.2 dpf. The number of fish analyzed by confocal microscopy is listed on the top. Refer Figure 2B for valve morphology classification. (**D**) Selumetinib treatment can decrease the edema ratio in *rasa1a^tsu40^;rasa1b^tsu39^* double mutants at proper concentrations as indicated. The number of fish analyzed after genotyping is listed on the top. Fisher’s exact test, ****, p < 0.0001. (**E**) Selumetinib treatment restores the LV and LVV formation in *rasa1a^tsu40^;rasa1b^tsu36^* mutants at 4 dpf. Prox1a immunostaining was used to label the valve-forming LECs. Arrowheads and arrows indicate the LV and FCLV-PHS LVV, respectively. The ratio of embryos with exhibited pattern of the Prox1a expression (cyan) is indicated. Lateral views, anterior to the left. Scale bars, 20 μm. (**F**) Statistical analysis of the high-level Prox1a-positive valve-forming LECs in siblings with DMSO (n = 4), *rasa1a^tsu40^;rasa1b^tsu36^* mutants with DMSO (n=4), and *rasa1a^tsu40^;rasa1b^tsu36^* mutants with 100 μM Selumetinb (n = 4) in (E). Unpaired *t* test, ns, no statistical significance; ***, p < 0.001; ****, p < 0.0001.

## Discussion

Activation of MAPK/Erk signaling by Vegfc-Vegfr3 is critical to induce lymphatic endothelial cells (LECs) fate and therefore promoting lymphatic vessels formation, whereas excessive MAPK/Erk activation always leads to lymphatic abnormalities seen in related lymphatic disease (Baek et al., 2019; Deng et al., 2013; Li et al., 2019; Pandit et al., 2007). However, loss of both *Spred1* and *Spred2* in mice, two genes that act to inhibit Vegfr3 signaling-activated Erk1/2, causes not only LEC overgrowth, but also embryonic edema and blood filling of lymphatic vessels (Taniguchi et al., 2007), implying that MAPK/Erk signaling may also be important for blood-lymph separation, and possibly for LVV formation.

In this study, we discover that inhibition of the Ras-Raf-Mek-Erk cascade by Efnb2-Ephb4- Rasa1 signaling is required for the fate specification of valve-forming LECs during the formation of the zebrafish lymphatic system. Lack of *efnb2, ephb4b,* or *rasa1* leads to the increased Erk1/2 activation and defective LV or LVV formation, whereas simultaneous loss of Erk1 and Erk2 causes hyperplasia of the valve-forming LECs at the expense of some LECs (Figure 8). Moreover, the abnormal lymphatic valves due to *ephb4b* or *rasa1* deficiency can be partially rescued by pharmacological inhibition of Mek-Erk activation. Collectively, our findings demonstrate that the MAPK/Erk cascade needs to be repressed during the valve-forming LECs fate determination, which differs from its positive role in the lymphatic endothelial cells (LECs) specification. And the inhibition of MAPK/Erk signaling during lymphatic valve formation is fulfilled by Efnb2-Ephb4b-activated Rasa1, uncovering an important biological role of the crosstalk between Efnb2-Ephb4b and MAPK/Erk signaling under physiological conditions.

**Figure 8.**
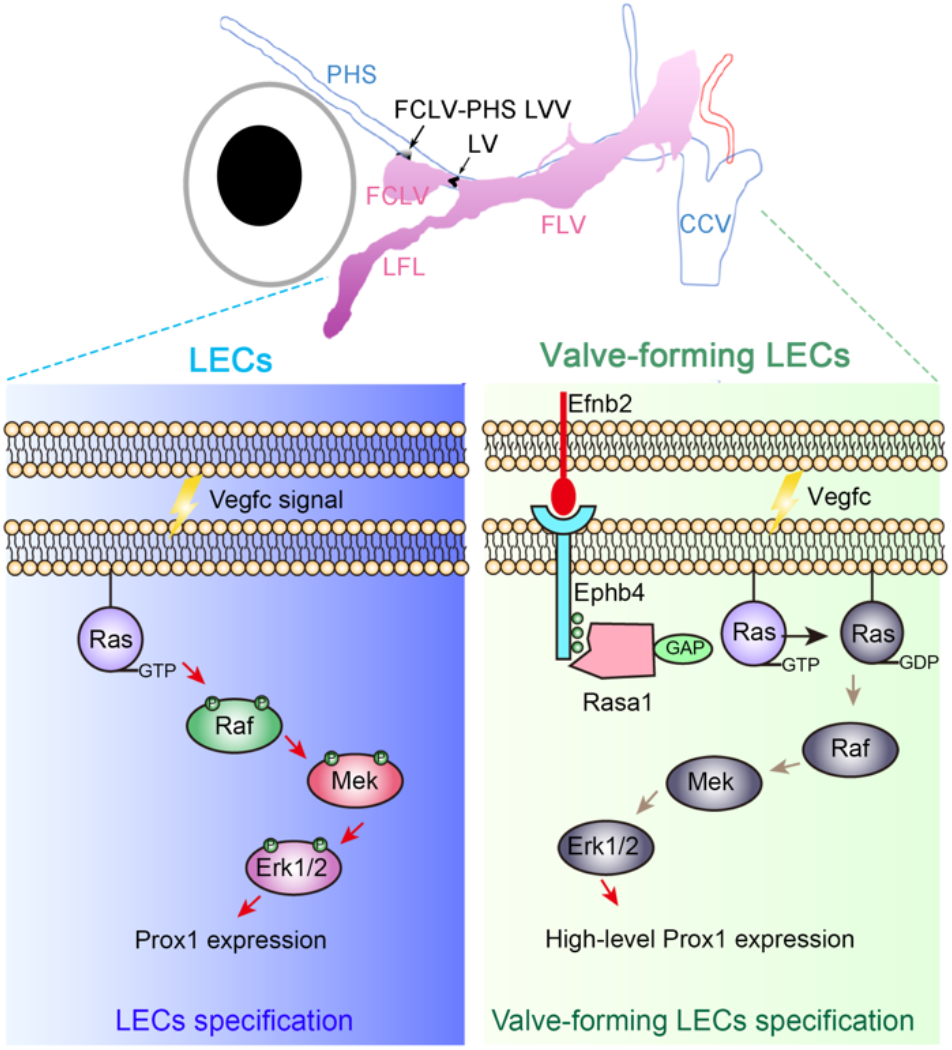
Working model of Efnb2-Ephb4-Rasa1 in valve-forming LECs specification. In lymphatic endothelial cells (LECs), Vegfc signaling stimulates Ras-Raf-Mek-Erk pathway for Prox1 expression and subsequently LECs specification. In valve-forming LECs, Efnb2-Ephb4 signaling recruits Rasa1, a Ras GTPase activating protein, to inactivate Ras-GTP to Ras-GDP, and therefore inhibiting Ras-Raf-Mek-Erk pathway specifically. This kind of inhibition somehow allows a high expression level of Prox1 that is required for the valve-forming LECs specification.

Actually, Vegfc-Vegfr3 also activates PI3K-Akt signaling in lymphatic endothelial cells (Deng, Zhang, & Simons, 2015). PI3K-Akt signaling has been shown to stimulate de novo lymphatic valve growth in mice, potentially by inactivating Foxo1, a key negative regulator to inhibit the expression of valve-forming genes, including *Foxc2*, *Gata2*, and *Prox1* (Scallan et al., 2021). Considering that, in lymphatic endothelial cells, PI3K-Akt inhibits MAPK/Erk signaling via Akt1-dependent phosphorylation of Raf1 on Ser259 (Deng et al., 2013; Ren et al., 2010), Akt1-Raf1 crosstalk might also be involved in valve-forming lymphatic endothelial fate specification by controlling Erk1/2 activation. Combined with our findings, we believe that the inhibition of MAPK/Erk is vital for the lymphatic valve formation, and might be achieved by synergistic interaction of several signaling pathways. However, the molecular mechanism of MAPK/Erk to repress the expression of valve-forming genes remains elusive and needs further investigation.

Although Efnb2-Ephb4 signaling has been implied in the regulation of lymphatic valve maturation during late embryonic and early postnatal development in mice (Mäkinen et al., 2005; G. Zhang et al., 2015), this is the first time to explore its vital roles in the cell fate determination of lymphatic valve progenitor cells. Here, we suppose that the Efnb2-Ephb4-Rasa1 forward signaling pathway provides a critical signal to stimulate high-level Prox1 expression in valve-forming progenitor cells. A series of increasing whole exome sequencing studies have revealed that the EFNB2-EPHB4-RASA1 signaling axis is important for human vascular diseases such as capillary malformation-arteriovenous malformation (CM-AVMs), vein of Galen malformation (VOGM) and so on (Amyere et al., 2017; Eerola et al., 2003; Revencu et al., 2008; J. Yu, Streicher, Medne, Krantz, & Yan, 2017; Zeng et al., 2019). In addition to the malformed blood vasculature, lymphatic abnormalities can also be observed in some CM-AVM and VOGM patients, and *EPHB4* mutations are found in patients with lymphatic disorder central conducting lymphatic anomaly (CCLA) (Burrows et al., 2013; Li et al., 2018). These findings indicate a complex regulation of blood and lymphatic vessels under the control of EFNB2-EPHB4-RASA1 signaling. Here our findings not only illustrate a possible mechanism underlying the lymphatic abnormalities in such diseases, but also establish several zebrafish genetic disease models, such as *ephb4b^tsu25^*, *efnb2a^tsu41^*;*efnb2b^tsu42^* and *rasa1a^tsu38^;rasa1b^tsu39^* to further understand the pathogenesis of human lymphedema and evaluate potential drugs. More importantly, we found that treatment with MEK inhibitors significantly improves lymphedema phenotype and restores the lymphatic valve formation in *ephb4b^tsu25^* or *rasa1a^tsu40^;rasa1b^tsu39^* mutant larvae, providing a potential treatment strategy for valve-deficient disorders that currently lack specific molecular treatments.

We noted that LVs and LVVs almost fail to form in *efnb2a^tsu41^*;*efnb2b^tsu42^* or *rasa1a^tsu38^;rasa1b^tsu39^* double mutants, but partially form in *ephb4b^tsu25^* mutant larvae, which is consistent with the differences in the numbers of the valve-forming LECs, suggesting that the severity of lymphatic valve defects is not exactly the same in these mutants. Considering the fact that *efnb2a/2b* expression occurs in both FCLV and LFL, whereas *ephb4b* is only expressed in LFL LECs (Figure 6C and S4B), we suspect that both FCLV and LFL might contribute to the valve-forming LECs fate specification. In addition, Efnb2 ligand is able to activate other Eph receptors such as Epha4, Ephb1 or Ephb3 in certain context (Gucciardo, Sugiyama, & Lehti, 2014; Kullander & Klein, 2002; Murai & Pasquale, 2003; Noren & Pasquale, 2004), which may be expressed in FCLV LECs and thus regulate lymphatic valve formation. These possibilities need to be investigated in future studies.

## Materials and Methods

### Zebrafish and Maintenance

Zebrafish were raised and maintained with ethical approval from the Animal Care and Use Committee of Tsinghua University. The generation of mutants or transgenic lines is described below. Most of the lines were crossed with *Casper* mutant (White et al., 2008) to obtain non-pigment embryos. Otherwise, 0.003% 1-phenyl-2-thiourea (PTU, Sigma, P7629) was used to inhibit pigment formation. *Tg(gata2:EGFP)^la3^*, *Tg(gata1:DsRed)^sd2^* (Traver et al., 2003), *Tg(flk:EGFP)^s843^* (Jin, Beis, Mitchell, Chen, & Stainier, 2005), and *Tg(flk:mCherry)* (Xia et al., 2013) are described before.

### Transgenic Fish Generation

The transgenic fish with *lyve1b* promoter *Tg(lyve1b:TopzaYFP)^tsu47tg^* and *Tg(lyve1b:DsRed2)^tsu48tg^* were generated following the protocol published by (Okuda et al., 2012). The corresponding 5.2 kb *lyve1b* promoter was amplified and inserted into a plasmid vector containing Tol2 elements. Briefly, 20 pg of plasmid and 200 pg of Tol2 transposase mRNA were co-injected into 1-cell embryos with Tuebingen background and founder fish was identified by crossing with wild-type adult fish. See Table S2 for primer information.

### Generation of Knockout Lines and Genotyping

For making knockout lines, we injected Cas9 mRNA or protein (NEB, M0646T) with corresponding sgRNAs into 1-cell wild-type Tuebingen embryos. Cas9 mRNA was synthesized using the mMESSAGE mMACHINE T7 Transcription kit (ThermoFisher, AM1344). gRNA was synthesized using the MEGASCRIPT T7 Kit (ThermoFisher, AM1334). The Cas9 mRNA used for injection was about 200 pg, or Cas9 protein was 1-3 fmol, and sgRNAs were 100-400 pg per embryo. Founder fish and F1 adults with mutations were identified by sequencing. The F1 or F2 adult fish were then mated with *Casper* mutant. See Table S1 for additional allele information and Table S2 for mutant identification.

### Pericardial Injection for Lymphangiography

2,000 kDa Dextran-Fluorescein (Invitrogen, D7137) was used in this study. Anesthetized embryos at 3-3.2 dpf were placed laterally on an agarose plate with a single hole. For embryo pericardial lymphangiography, 1 nl of 10 mg/ml Dextran-Fluorescein was injected into the pericardial region and fluorescence was observed under an Olympus MVX10 stereomicroscope at 1 hour post injection (hpi) or 24 hpi.

### Whole Mount Immunostaining

Fish with mutations and transgenic background at 3.2 or 4 dpf were fixed 1 to 2 days at 4℃ with 4% paraformaldehyde (Sigma, P6148) in PBS. They were then transferred to 100% methanol and stored at -20℃ for at least 1 day. For detection of pERK and Prox1, the embryos were treated with 1 mM EDTA (pH 8.0) at 90℃ for 10 min and then blocked in 2% BSA for 2 hours. Rabbit anti-Phospho-ERK(Thr202/Tyr204) (1/200; CST, 4370), rabbit anti-Prox1 (1/200; GeneTex, GTX128354), chicken anti-GFP (1/200; Abcam, ab13970), and mouse anti-mCherry (1/100; EASYBIO, BE2026) were used for primary incubation. Goat anti-chicken IgY-Alexa Fluor 488 (1/400; Abcam, ab150169), goat anti-rabbit IgG-Alexa Fluor 647 (1/400; Jackson, 111-605-003), and goat anti-mouse IgG-TRITC (1/400; Jackson, 115-545-003) were used for secondary incubation. Embryos were mounted on glass slides with proper height adhesion tape and covered by coverslips.

### Whole Mount Fluorescence *In Situ* Hybridization (FISH)

Whole mount fluorescence *in situ* hybridization was carried out as described previously (He, Mo, Chen, & Luo, 2020), but without the process of peeling skin. TSA Plus Cyanine 5 Kit (Akoya Biosciences, NEL745001KT) was used. Embryos were treated with 1 mM EDTA (pH 8.0) at 90℃ for 10 min before prehybridization. Eyes were removed for mounting and imaging after FISH. DIG-labeled antisense or sense probes were synthesized using T7 or T3 RNA polymerase (Roche, 10588423 and 11031171001) and Dig RNA Labeling Mix (Roche, 11277073910). Sometimes, Prox1 antibody and 1/400 goat anti-rabbit IgG-Alexa Fluor 488 (Jackson, 111-545-003) were used for immunostaining after FISH. PCR products were used as templates, and primers used for generating probe templates are listed in Table S2.

### Imaging and Image Processing

To observe the lymphatic vessels or valve structure, embryos or larvae were anesthetized with 1 or 0.2 mg/ml Tricaine (Sigma, A5040) and embedded in 1% low melting agarose/Holtfreter’s water in 35-mm glass bottom culture dishes. Imaging was carried out on a Perkin Elmer Spinning Disk confocal or Dragonfly Spinning Disk confocal microscope (Andor) using a 20x objective. For FISH or immunostaining, a 40x oil objective was used. Blood and lymph autofluorescence can be detected by the 561 channel. Embryos with unknown genotypes were imaged first, then lysed with 30 μl 50 mM NaOH at 95℃ for 20 min and genotyped by appropriate methods (See Table S2). The images were then viewed and processed by Imaris software 9.1. Single plane or 3D images were generated by Snapshot and then processed by Adobe Photoshop 2020. Movies were generated using Imaris 9.1, imported into Adobe Premiere Pro 2020 for labeling, and exported as .mp4 files.

### Flow Cytometry

Single cell suspension of 3.2 dpf embryos was generated by treating with 50 μl TrypLE (ThermoFisher, 12604013) per embryo at 28.5℃ for 2-3 hours with pipetting every 20 min. The cell suspension was then filtered through a 35-μm nylon mesh (Falcon, 352235), centrifuged at 1000 *g* for 5 min at 4℃, and then resuspended in 5% fetal bovine serum (FBS) in PBS. DAPI (1 mg/ml; Enzo Life Sciences, BML-AP402-0010) was added to exclude the dead cells. FSC and SSC were used to exclude debris. Flow cytometry was performed on a MoFlo XDP (Beckman Coulter). The cells were collected in 5% FBS in PBS, and then concentrated by centrifugation. The concentrated cell suspension was applied on the glass slide for pERK immunostaining. The slides were observed on a Dragonfly confocal microscope (Andor) and analyzed by Imaris.

### Small Molecules Treatment

For inhibitor treatment in *ephb4b^tsu25^* mutants, embryos were raised to 2 dpf at 28.5℃ and then transferred into 6-well plates with 30 embryos per well. We added 3 ml 25% Holtfreter’s water with proper concentrations of small molecules or an equal amount of DMSO, and then the embryos were raised to 4 dpf at 30.5℃ to observe pericardial edema. Rapamycin (Solaribio, R8140-25), BEZ235 (Topscience, T2235), and MEK inhibitors Selumetinib (Topscience, T6218), Cobimetinib (Topscience, T3623), Trametinib (Topscience, T2125) and U0126-EtOH (Topscience, T6223) were used. For inhibitor treatment of *rasa1* mutants, 10 or 100 μM Selumetinib were used from 2.4 dpf to 4 dpf in 100% Holtfreter’s water at 30.5℃. Inhibitor effects were measured by comparing the pericardial edema in inhibitor treatment groups with pericardial edema in DMSO control groups.

### Statistical Analysis

Statistics analyses were performed in GraphPad Prism 8.2.1 and the mean ± SD was calculated. The unpaired student’s *t*-test, chi-squared test and Fisher’s exact test were used to calculate significance. The sample size (n) and *P* value for each experimental group are described in corresponding figure legends.

## Acknowledgments

We thank Dr. Yiyue Zhang for providing the *Tg(gata2:GFP)^la3^* fish, Dr. Xinfeng Liu and Dr. Zheng Jiang for providing the *efnb2a* and *efnb2b* F1 fish, and Dr. Bo Zhang for providing the Cas9 reagents. We also thank Dr. Yeqi Wang for providing the Prox1 antibody. We are grateful to State Key Laboratory of Membrane Biology and the Image Core Facility of Protein Technology Center of Tsinghua University for imaging assistance.

## Author contributions

YPM performed most experiments. TL generated the *erk1* and *erk2* mutants. JFZ generated the *ephb4b* mutants. YPM, AMM and SJJ designed the study and prepared the manuscript.

## Data and materials availability

All data are available in the main text or the supplementary materials.

## Source data Files

**Figure 3-source data 1**

**Ratio of embryo with pericardial edema.**

**Figure 6-source data 1**

**List of intensity mean of gata2:EGFP and lyve1b:DsRed2 positive endothelial cells.**

**Figure 7-source data 1**

**Inhibitor treatments for *ephb4b* mutant.**

**Video 1.**

**Slow red blood cells flow in lymphatic vessels in *ephb4b^tsu25^* mutants.** Solid lines indicated the blood in a lymphatic vessel. The RBCs in non-blood vessels or blood-vessels are indicated by white or yellow arrows, respectively.

**Video 2.**

**FCLV-PHS LVV can block RBCs flow from PHS into facial lymphatic vessels.** In *ephb4b^tsu25^* mutants, the absence of FCLV-PHS LVV allows the RBCs enter the facial lymphatic vessels.

**Video 3.**

**Lack of valve specification in *efnb2a^tsu41^;efnb2b^tsu42^* mutant. at 3.2 dpf by Prox1a immunostaining.** Immunofluorescence results of Prox1a (cyan) in siblings and *efnb2a^tsu41^;efnb2b^tsu42^* mutants at 3.2 dpf.

---

**Figure 1-figure supplement 1.**
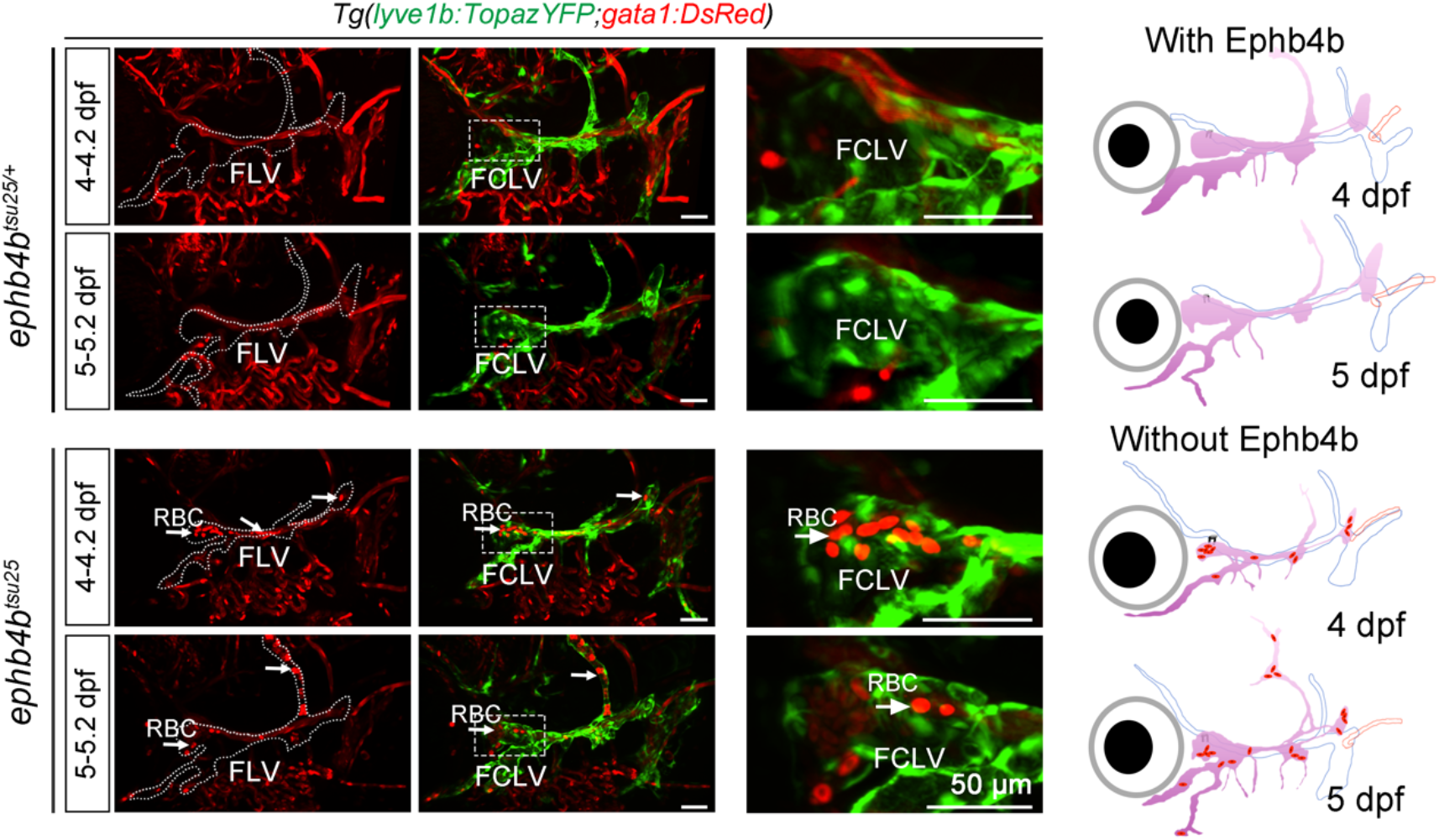
Red blood cell accumulation in *ephb4b^tsu25^* lymphatic vessels. *gata1:DsRed* labeled RBCs enter the *lyve1b:TopazYFP* labeled facial lymphatic vessels (green) at 4-4.2 dpf and 5-5.2 dpf in *ephb4b^tsu25^* mutants (arrows). Dotted lines indicate the facial lymphatic vessels (FLVs). The third panels are enlarged regions representing the facial collecting lymphatic vessels (FCLVs). Schematic diagrams in the right panels indicate the accumulation of RBCs in FLVs when Ephb4b is lost. Lateral views, anterior to the left, dorsal to the top. Scale bars, 50 μm.

**Figure 1-figure supplement 2.**
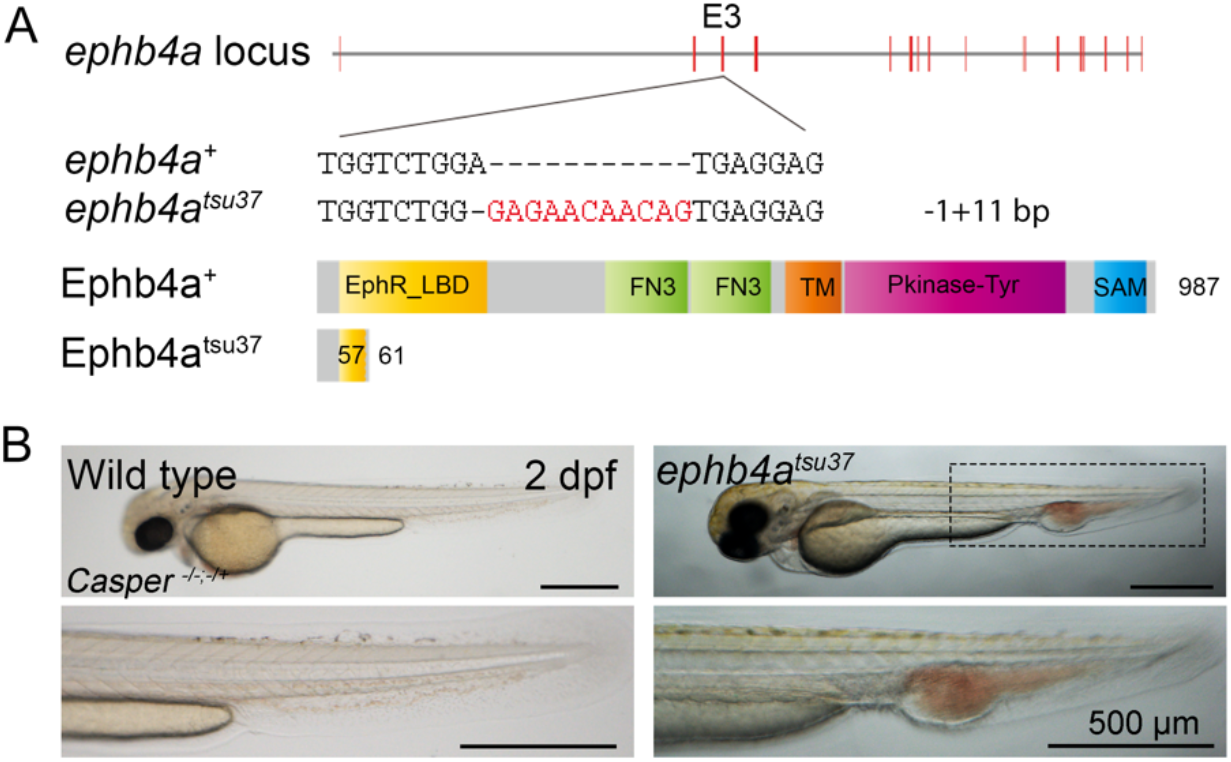
The *ephb4a^tsu37^* mutants exhibit caudal blood vessel malformation. (A) Generation of the *ephb4a* mutant using CRISPR/Cas9 technology with a target site at exon 3. The *ephb4a^tsu37^* mutant bears a 1-bp deletion and 11-bp insertion with a protein length of 61 aa. The amino acid sequence changes after 57 aa. The *ephb4a^tsu37^* mutant protein only has a partial ligand binding domain (LBD). (B) *ephb4a^tsu37^* mutants exhibit abnormal blood vessel formation and blood accumulation in the tail region at 2 dpf. Scale bars, 500 μm.

**Figure 2-figure supplement 1.**
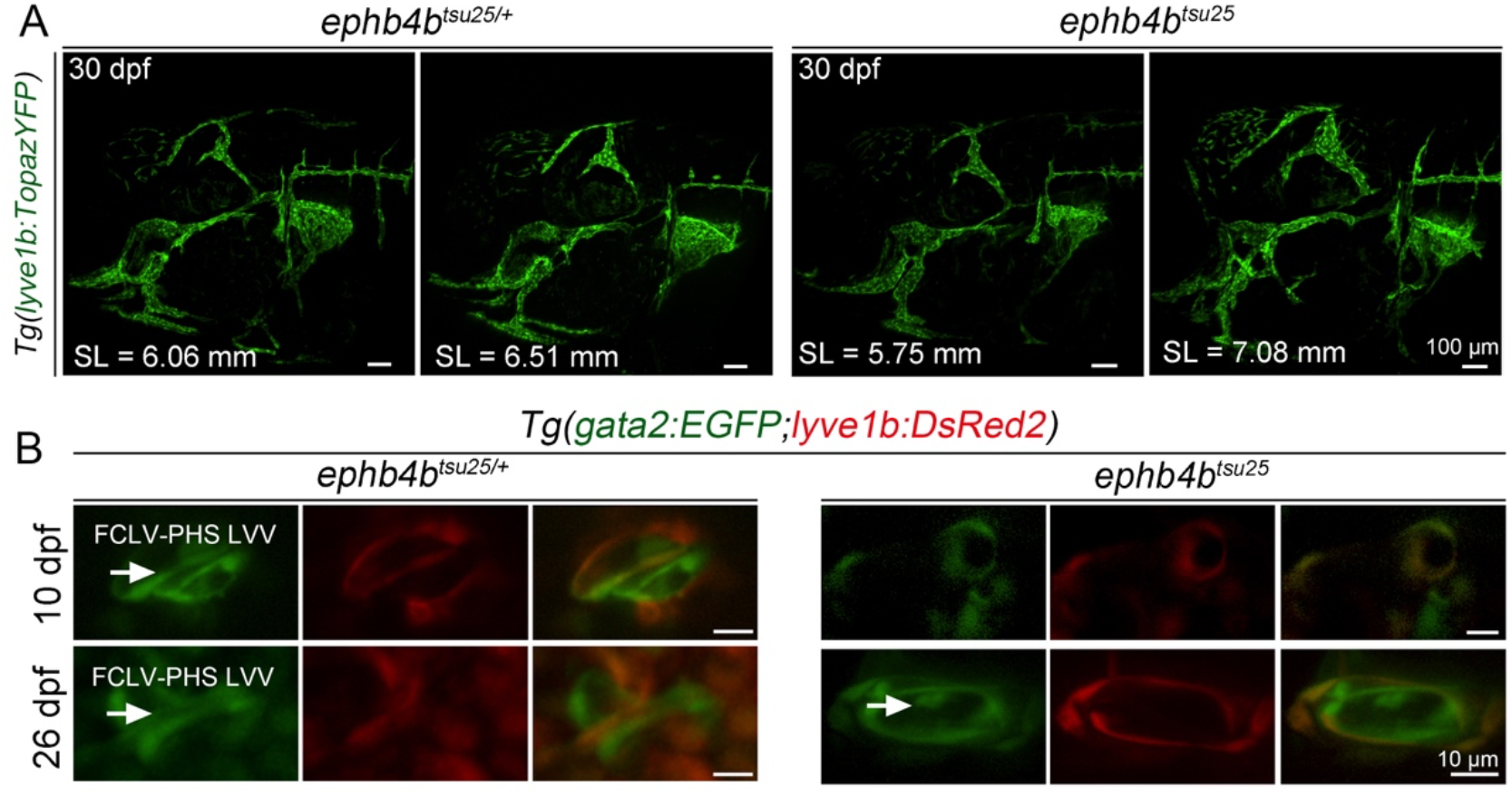
*ephb4b* is essential for lymphatic valve but not lymphatic vessel formation. (A) Formation of the lymphatic vasculature is normal in *ephb4b^tsu25^* mutants. Sibling and *ephb4b^tsu25^* mutant larvae in the *Tg(lyve1b:TopazYFP)* background were raised to 30 dpf. The *lyve1b:TopazYFP*-positive lymphatic vessels in the head region developed normally. SL, standard length. Lateral views, anterior to the left, dorsal to the top. Scale bars, 100 μm. (B) Defective formation of the FCLV-PHS LVVs in *ephb4b^tsu25^* mutants. Sibling and *ephb4b^tsu25^* mutant larvae in the *Tg(gata2:EGFP;lyve1b:DsRed2)* background were raised to 10 dpf and 26 dpf. *gata2:EGFP* labeled valve structure is defective in the mutant larvae. Scale bars, 10 μm.

**Figure 4-figure supplement 1.**
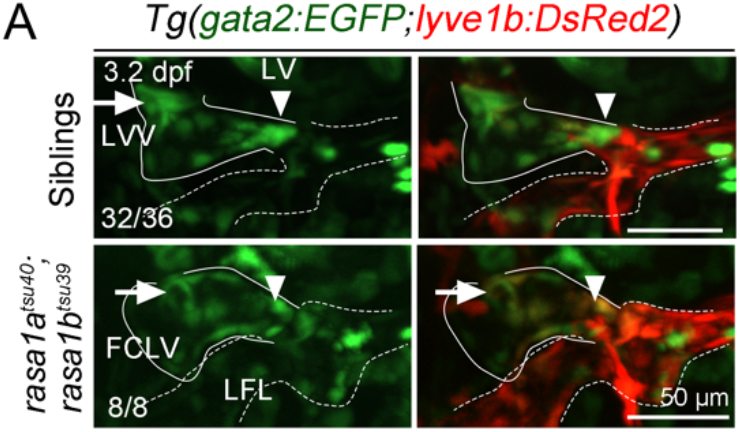
No valve structure formation in *rasa1a^tsu40^;rasa1b^tsu39^* double mutants. No valve structure formation in *rasa1a^tsu40^;rasa1b^tsu39^* double mutants at 3.2 dpf. The FCLV and lateral facial lymphatic vessel (LFL) in mutants are marked by solid lines and dotted lines respectively. LVs and FCLV-PHS LVVs are indicated by arrowheads and arrows, respectively. The ratio of embryos with exhibited valve structure is indicated. Siblings were defined as neither *rasa1a^tsu40^* mutant nor *rasa1b^tsu39^* mutants. Scale bars, 50 μm.

**Figure 6-figure supplement 1.**
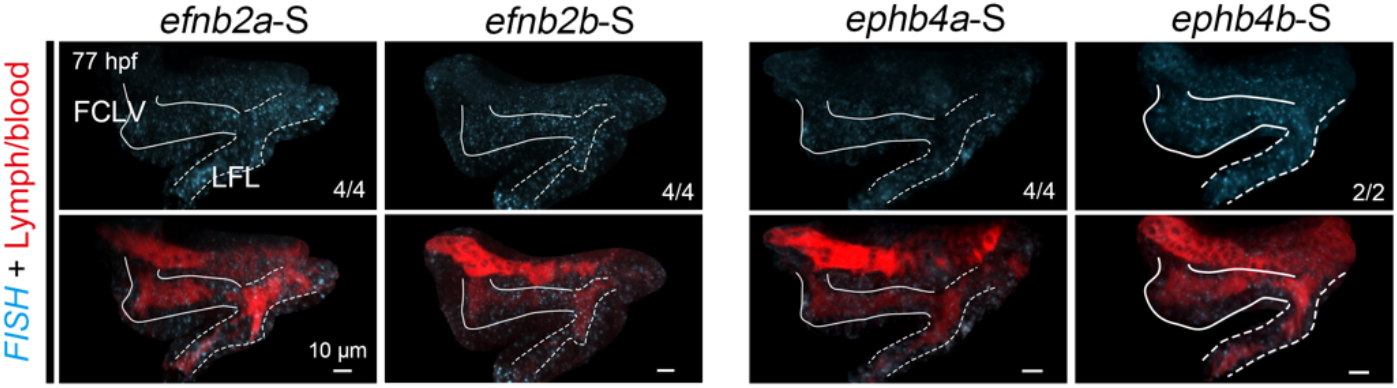
FISH staining results using sense probes as negative controls. Fluorescence *in situ* hybridization results for 77 hpf embryos using *efnb2a*, *efnb2b*, *ephb4a,* and *ephb4b* sense probes as control. Solid and dotted lines indicate FCLV and LFL, respectively. The ratio in the right corner indicates the number of embryos with the observed pattern to total embryos. Lateral-dorsal views, anterior to the left, dorsal to the top. Scale bars, 10 μm.

**Figure 7-figure supplement 1.**
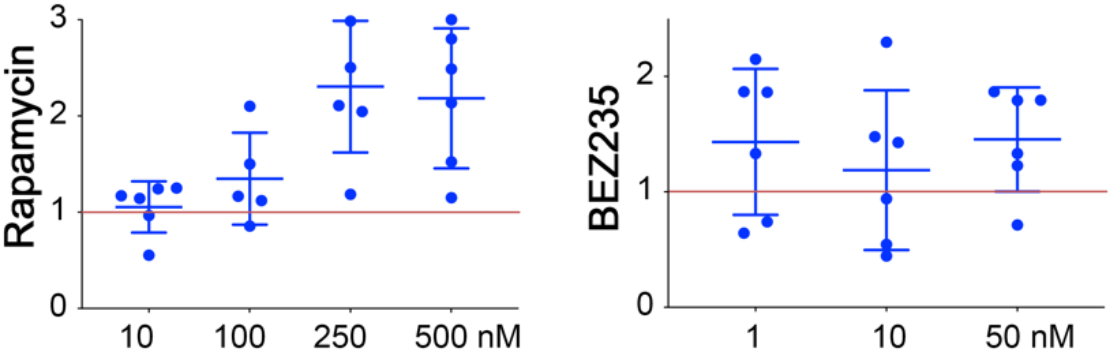
mTOR inhibitors cannot rescue the pericardial edema in *ephb4b^tsu25^* mutants. Related to Figure 7. mTOR inhibitor treatments cannot rescue the edema phenotype in *ephb4b^tsu25^* mutants. Rapamycin and BEZ235 were used at different concentrations as indicated. The vertical axis represents rescue effect value, and samples below the red lines indicate rescue effects.

**Table S1.**
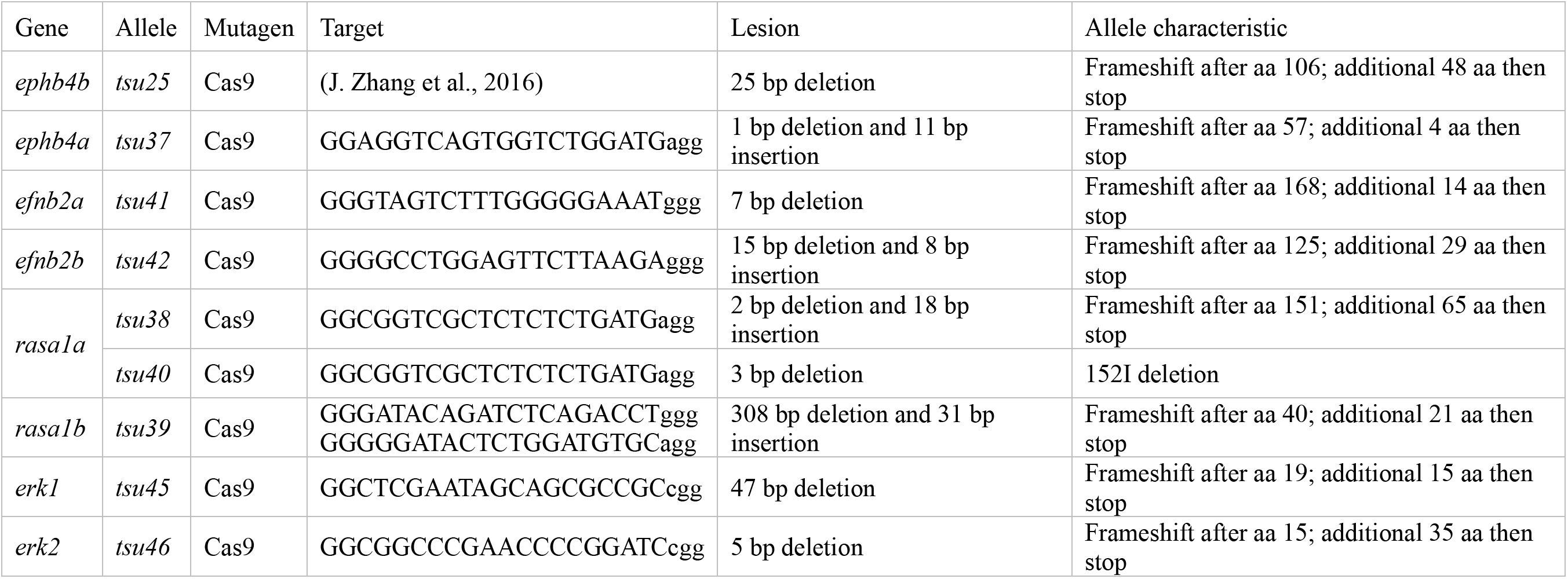
Zebrafish mutant lines used in this study.

**Table S2.**
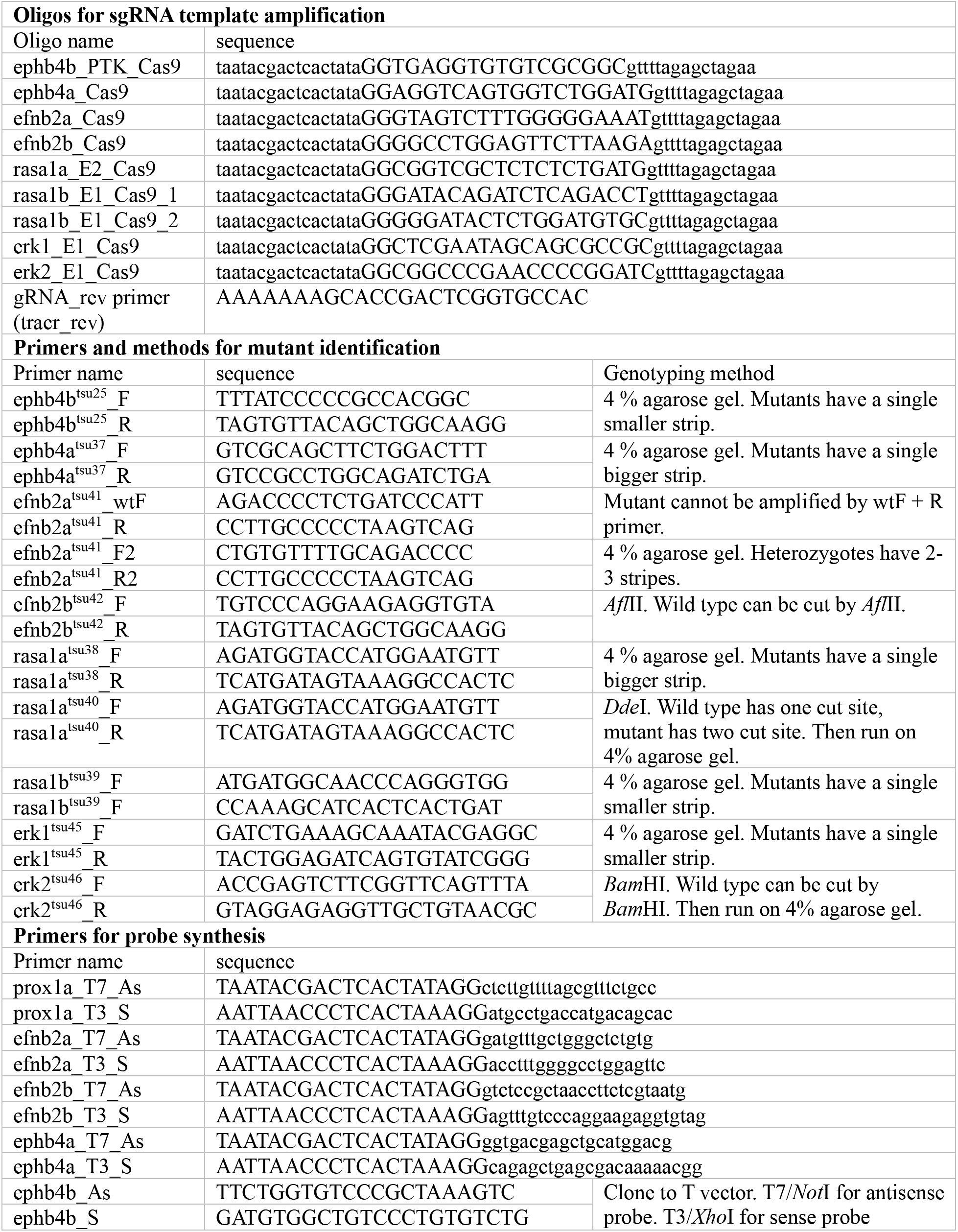
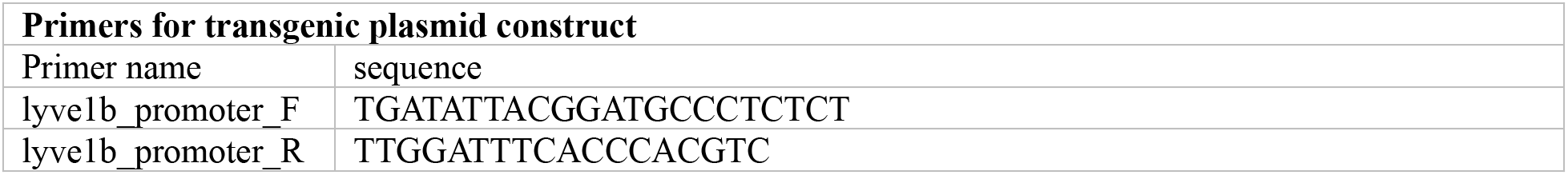
Oligonucleotides used in this study.

